# Type I interferon signaling promotes mucosal inflammation in murine models of colitis

**DOI:** 10.64898/2026.03.02.709022

**Authors:** Jianyi Yin, Rodrigo Galicia Pereyra, Luis Sifuentes-Dominguez, Emre E Turer, Ezra Burstein

## Abstract

Type I interferons (IFN-Is) play a critical role in innate immunity, modulating the host response. While dysregulated IFN-I signaling has been implicated in autoimmune and infectious disorders, its role in inflammatory bowel disease (IBD) remains unclear. In this study, we extensively assessed the function of IFN-I signaling in human IBD and murine models of colitis. Expression of IFN-I signature genes was elevated in patients with active ulcerative colitis as well as multiple murine models of colitis. Single cell RNA sequencing revealed that upregulated IFN-I signature genes were enriched in myeloid cells, which exhibited increased expression of IFN receptors during mucosal inflammation. Mice carrying gain-of-function alleles of *Ifnar1*, a subunit of IFN-I receptor, showed heightened IFN-I signaling and altered colonic immune homeostasis at baseline, and were more susceptible to experimental colitis. In contrast, postnatal inhibition of IFNAR1, using either an inducible transgenic mouse model or an anti-IFNAR1 blocking antibody, protected against experimental colitis. Taken together, our findings reveal a previously under-recognized pathogenic role of IFN-I in IBD and provide a rationale for therapeutic intervention targeting this pathway.

## INTRODUCTION

Interferons (IFNs) are structurally related cytokines that play essential roles in innate immunity (1). Organized into three distinct families, all IFNs are capable of inducing interferon-stimulated genes (ISGs) via Janus Kinase/Signal Transducer and Activator of Transcription (JAK/STAT) signal transduction pathways (2). Each IFN family signals through distinct receptors, and their biological functions do not fully overlap. Type I IFNs (IFN-Is) include 17 closely related proteins in humans (13 IFN-α family members, and IFN-β, IFN-ε, IFN-κ, and IFN-ω) that share a common receptor complex containing two subunits encoded by the *IFNAR1* and *IFNAR2* genes. IFN-Is are best known for their antiviral and immunomodulatory properties and act broadly on many cell types (3). Type II IFN (IFN-II) consists solely of IFN-γ, which binds a unique receptor complex encoded by the *IFNGR1* and *IFNGR2 genes*, exerting predominantly pro-inflammatory effects on immune cells (4). Finally, type III IFNs (IFN-IIIs) include four members (IFN-λ1/IL-29, IFN-λ2/IL-28A, IFN-λ3/IL-28B, and IFN-λ4) that bind to a receptor complex encoded by the *IFNLR1* and *IL10RB* genes. IFN-IIIs share functional similarities with IFN-Is, but their biological effects are largely confined to mucosal surfaces due to restricted expression of their receptors in specific cell types (5).

Globally, IFNs are essential for antiviral immunity, but dysregulated IFN responses have been linked to autoimmune disorders such as systemic lupus erythematosus (SLE) (6), or susceptibility to mycobacterial or viral infection (7). Recently, evidence has emerged linking an elevated IFN tone to the increased susceptibility for celiac disease in patients with Down syndrome (8). However, participation of IFNs in other gastrointestinal (GI) inflammatory disorders, such as inflammatory bowel disease (IBD), is less well studied. IBD is a chronic inflammatory disorder of the GI tract caused by a complex interaction between genetic factors and environmental triggers (9). Phenotypically, IBD is classified into two main subtypes: ulcerative colitis (UC), affecting the mucosa of the large intestine, and Crohn’s disease (CD), a transmural granulomatous inflammatory process that may affect any segment of the GI tract.

Impaired epithelial barrier function and dysregulated mucosal immune response are universally observed in IBD and are associated with abnormal expression of multiple cytokines, including IFNs (10, 11). While IFN-II is widely recognized as a contributor to IBD pathophysiology, the role of IFN-I in IBD is not well understood (4, 11). On the one hand, IFN-I can have anti-inflammatory effects and, in fact, IFN-β has been a first-line treatment for relapsing forms of multiple sclerosis (12). Nonetheless, when systemic IFN-I therapy was evaluated in IBD clinical trials, the results demonstrated no significant therapeutic efficacy (13). Moreover, *de novo* colitis or exacerbation of pre-existing IBD has been reported in patients receiving IFN-I for non-IBD conditions (14–16). Furthermore, recent studies have identified pathologic activation of IFN-I signaling in specific genetic disorders associated with early-onset IBD (17, 18). Thus, despite anti-inflammatory effects of IFN-I in some clinical settings, the human data appears to favor a deleterious role for IFN-I in IBD.

Pre-clinical studies in mouse models also paint a confusing picture for the role of IFN-I in experimental colitis. In support of a detrimental role of IFN-I in intestinal inflammation, local delivery of IFN-β via genetically modified *Lactobacillus acidophilus* exacerbates dextran sulfate sodium (DSS)-induced colitis (19). In contrast, other studies have reported increased susceptibility to DSS-induced colitis in IFNAR1 knockout (KO) mice and a protective effect of systemic administration of IFN-β during DSS-induced colitis (20). Of note, the phenotype of global IFNAR1-KO mice has been variable in other studies. One study reported increased susceptibility to colitis at a higher dose of DSS, but the opposite phenotype at a low dose (21). In another study, global IFNAR1 knockout led to mild disease worsening by some parameters but not others, whereas concurrent IFNAR1/IFNLR1 deficiency caused severe sensitivity to DSS-induced colitis, which was linked to decreased epithelial restitution due to decreased expression of the epithelial growth factor amphiregulin (22).

Altogether, the state of knowledge pertaining to the role of IFN-I in IBD is non-conclusive, with conflicting human and experimental data. Moreover, given that the IFN-I signaling pathway includes a number of therapeutic targets already in clinical use and others under investigation (23, 24), further understanding of the contribution of this pathway to IBD pathophysiology may inform the development of new treatment strategies. This is the impetus behind the present study, where we extensively evaluated IFN-I signaling in human IBD and murine models of colitis. Our results highlight a previously under-recognized pathogenic role of IFN-I signaling and provide a rationale for targeting this pathway for therapeutic purposes in IBD.

## RESULTS

### IFN-I and IFN-II signaling are upregulated in active UC

First, we investigated gene expression changes associated with disease activity by performing bulk RNA sequencing (RNA-seq) on colonic biopsies obtained from non-IBD controls (n=10), UC patients with inactive disease (endoscopic Mayo score of 0 with histologic finding of inactive colitis, n=6), and UC patients with active disease (endoscopic Mayo score of 2-3 with histologic finding of active colitis, n=11). All biopsy specimens were derived from the same colonic segment (sigmoid colon). Patient specimens were obtained from the University of Texas Southwestern Medical Center (UT Southwestern) biorepository for the study of inflammatory GI disorders, and patient characteristics in each group are included in **Supplemental Table 1**. Compared to non-IBD controls, a total of 2,057 differentially expressed genes (DEGs) were significantly upregulated and 682 were downregulated in biopsies from patients with active UC **(Figure 1A)**. Principal component analysis based on these transcriptional signatures revealed that active UC patients clustered separately from inactive UC patients and non-IBD controls, with the latter two groups largely overlapping, suggesting that active mucosal inflammation is the main driver of these transcriptional changes **(Figure 1B)**. Gene ontology enrichment analysis of the upregulated DEGs showed that biologic processes related to responses to IFN-α (IFN-I), IFN-β (IFN-I), and IFN-II were significantly enriched, whereas altered response to IFN-III was not found **(Figure 1C)**. Additional pathway enrichment analyses using Molecular Signatures Database (MSigDB) and Reactome Pathway Database consistently identified IFN-I and IFN-II signaling among the most enriched pathways in active UC **(Supplemental Figure 1A-B)**. These RNA-seq findings were validated by quantitative reverse transcription PCR (qRT-PCR), which demonstrated consistent elevation of several ISGs in colonic biopsies from patients with active UC **(Figure 1D)**, as well as upregulation of inflammatory genes like *IL6* and *S100A8* **(Supplemental Figure 1C)**. In addition to UC, analysis of the IBD TaMMA database (25) showed that several ISGs were also significantly upregulated in colonic specimens from patients with CD compared with non-IBD controls **(Supplemental Figure 1D)**, further supporting a pronounced ISG signature during mucosal inflammation across IBD subtypes.

**Figure 1.**
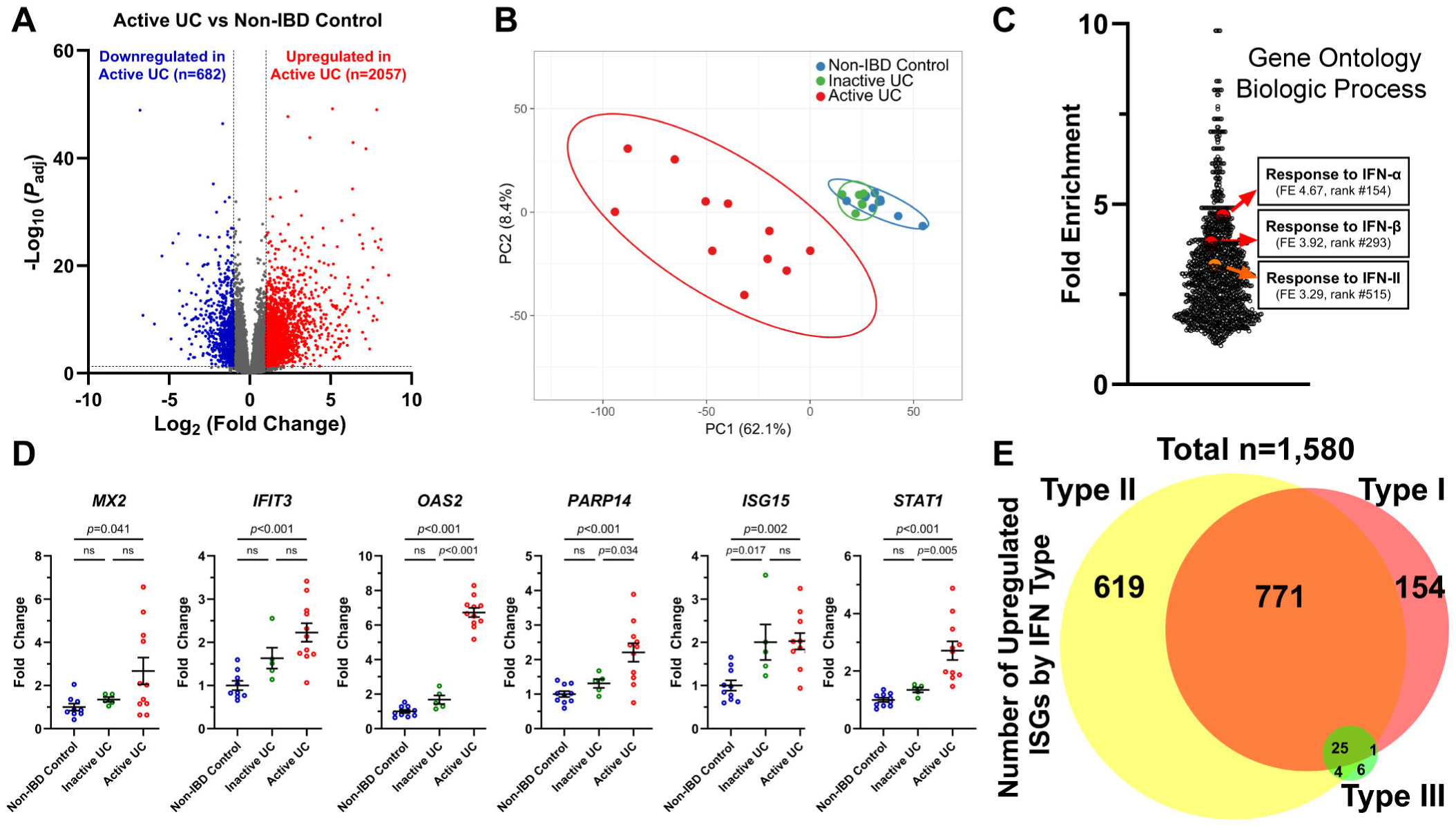
IFN-I signaling is activated in active ulcerative colitis (UC). **(A)** Volcano plot showing differentially expressed genes (DEGs, fold change >2, FDR-adjusted *p* <0.05) from bulk RNA sequencing analysis of colonic biopsies from active UC patients (n=11) vs non-IBD controls (n=10). **(B)** Principal component analysis of colonic biopsies from active UC patients, inactive UC patients, and non-IBD controls using the DEGs identified in (A). PC: principal component. **(C)** Gene ontology enrichment analysis of the upregulated DEGs identified in (A). FE: fold enrichment. **(D)** qRT-PCR validation of selected interferon stimulated genes (ISGs) in colonic biopsies. n=6-11 per group. ns: not significant. **(E)** Venn diagram of upregulated DEGs in (A) identified as ISGs and their IFN type according to the Interferome database.

Next, we sought to develop a more comprehensive view of the DEGs that belong to the ISG class. To accomplish this, upregulated DEGs were evaluated using the Interferome database, a publicly available resource for interpreting ISGs based on published transcriptomic studies (26). A total of 1,580 upregulated DEGs were identified as possible ISGs with evidence of transcriptional changes in response to IFNs in prior *in vivo* and *in vitro* studies **(Figure 1E)**. Interestingly, most upregulated ISGs were IFN-II-inducible genes (n=1,419 [89.8%]), followed by IFN-I-inducible genes (n=951 [60.2%]), and very few IFN-III-inducible genes were noted (n=36 [2.3%]). This analysis favored a substantial contribution of IFN-I and IFN-II pathways, but significant overlap was evident, as 50.4% of the upregulated ISGs (796 genes) could be induced by both IFN-I and IFN-II **(Figure 1E)**.

### IFN-I signature genes are upregulated in human IBD and murine colitis

Next, we overlapped the genes upregulated in active UC and annotated in the IFN-I response pathway across the three gene ontology databases described above (MSigDB, Reactome and Gene Ontology) and identified 54 ISGs to represent the IFN-I signature **(Figure 2A)**. Of note, a search in the Interferome database showed that nearly all (n=53 [98.1%]) of these 54 IFN-I signature genes are also inducible by IFN-II, whereas only a small subset (n=16 [29.6%]) are inducible by IFN-III **(Supplemental Table 2)**. Transcript levels of these 54 ISGs were consistently higher in colonic biopsies from patients with active UC than biopsies from non-IBD controls or patients with inactive UC, independent of IBD medications used at the time of specimen collection **(Figure 2B)**. Subsequently, we used immunofluorescence staining and image quantification to examine the protein expression of two representative ISGs in this group: ISG15, one of first identified and most well-known ISG, and STAT1, the key downstream transcription factor in IFN-I signaling, which itself is an ISG. Using fixed colonic biopsies from the same colonic segment (rectum) in a separate cohort of patients with active UC (n=5) and non-IBD controls (n=5) from our biorepository **(Supplemental Table 1)**, we observed that protein expression of both ISG15 and STAT1 was significantly upregulated in active UC **(Figure 2C-D)**. Interestingly, the immunofluorescence signals for both proteins were predominantly localized to lamina propria cells **(Figure 2C)**.

**Figure 2.**
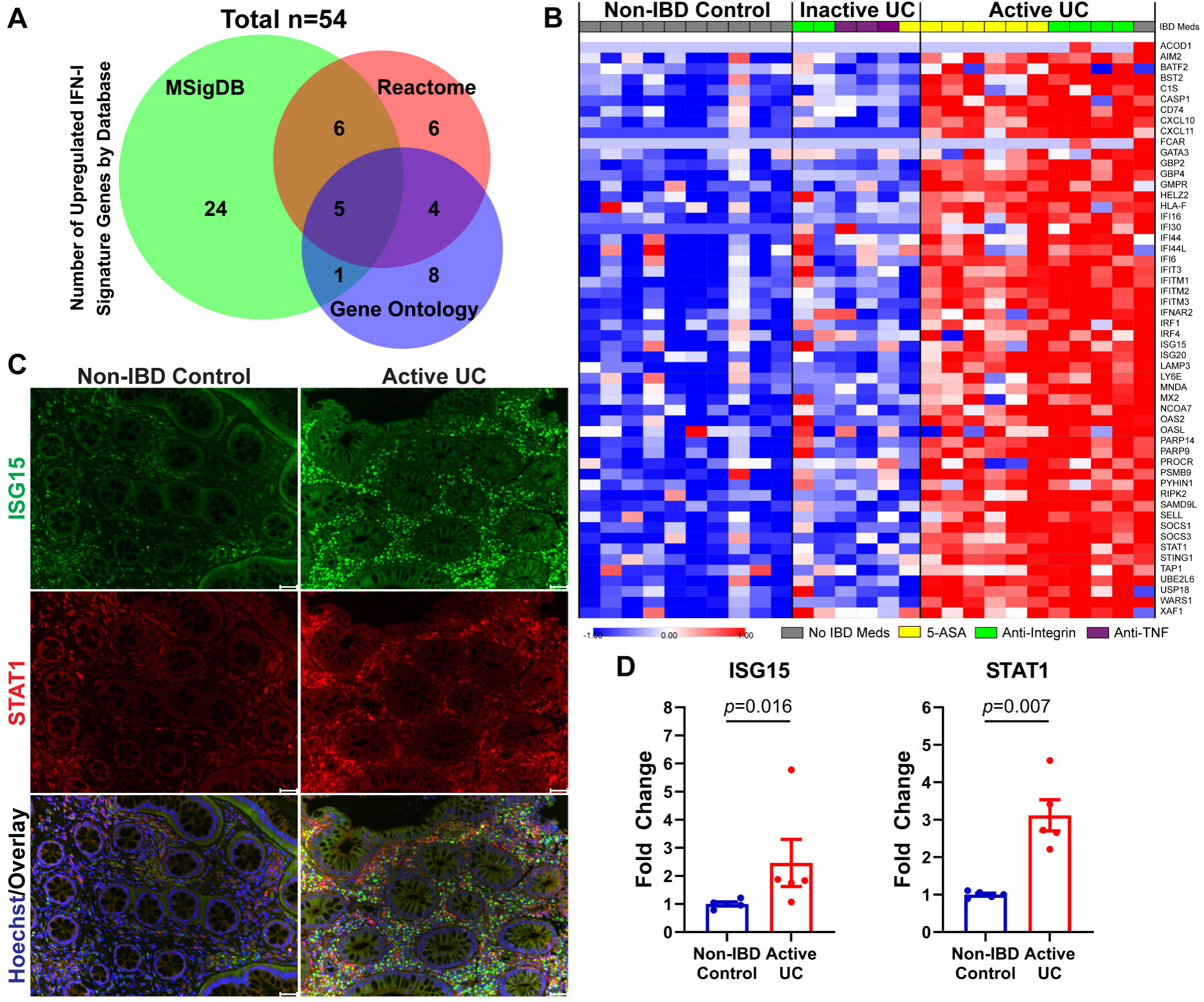
IFN-I signature genes are upregulated in active ulcerative colitis (UC). **(A)** Venn diagram of interferon-stimulated genes (ISGs) in response to IFN-I among the upregulated DEGs from Figure 1A, based on annotations from three pathway enrichment databases (MSigDB, Reactome, Gene Ontology). A total of 54 ISGs were identified as representative IFN-I signature genes. **(B)** Heatmap showing the expression of the 54 IFN-I signature genes in colonic biopsies from non-IBD controls (n=10), inactive UC patients (n=6), and active UC patients (n=11). 5-ASA: 5-aminosalicylic acid. TNF: tumor necrosis factor. **(C-D)** Representative immunofluorescence images of ISG15 and STAT1 in colonic biopsies and quantification of immunofluorescence signals. Intensities were normalized to the mean of non-IBD controls (set as 1). n=4-5 per group. Scale bar: 50 μm.

We then asked whether murine models of experimental colitis recapitulate the induction of IFN-I signaling and upregulation of IFN-I signature genes observed in human IBD. We re-analyzed a publicly available bulk RNA-seq dataset from a study of DSS-induced colitis (GSE131032) (27). Among the 54 IFN-I signature genes identified in our RNA-seq analysis in human UC biopsies, 48 were detected in this dataset and exhibited an overall increase in expression following DSS exposure **(Figure 3A)**. Using qRT-PCR, we validated the increased expression of selected ISGs (*Mx2*, *Ifi44*, *Oas2*, and *Isg15*) in our own cohort of mice with DSS-induced colitis **(Figure 3B)**. Similar changes were also noted in a piroxicam-accelerated enterocolitis model (28), in which enterocolitis develops upon piroxicam exposure in *Il10* knockout (IL10-KO) mice but not in wild-type (WT) littermate controls **(Figure 3C)**. Consistent with these transcriptional changes, immunofluorescence staining of colonic tissue coupled with quantitative image analysis demonstrated increased protein expression of ISG15 and STAT1 in both murine models **(Figure 3D–G)**. Similar to the observation in human UC biopsies, ISG15 and STAT1 signals in both murine models were predominantly present in lamina propria cells **(Figure 3D and F)**. Altogether, these studies indicate that induction of IFN-I signature genes is a common feature of colitis in human and murine models.

**Figure 3.**
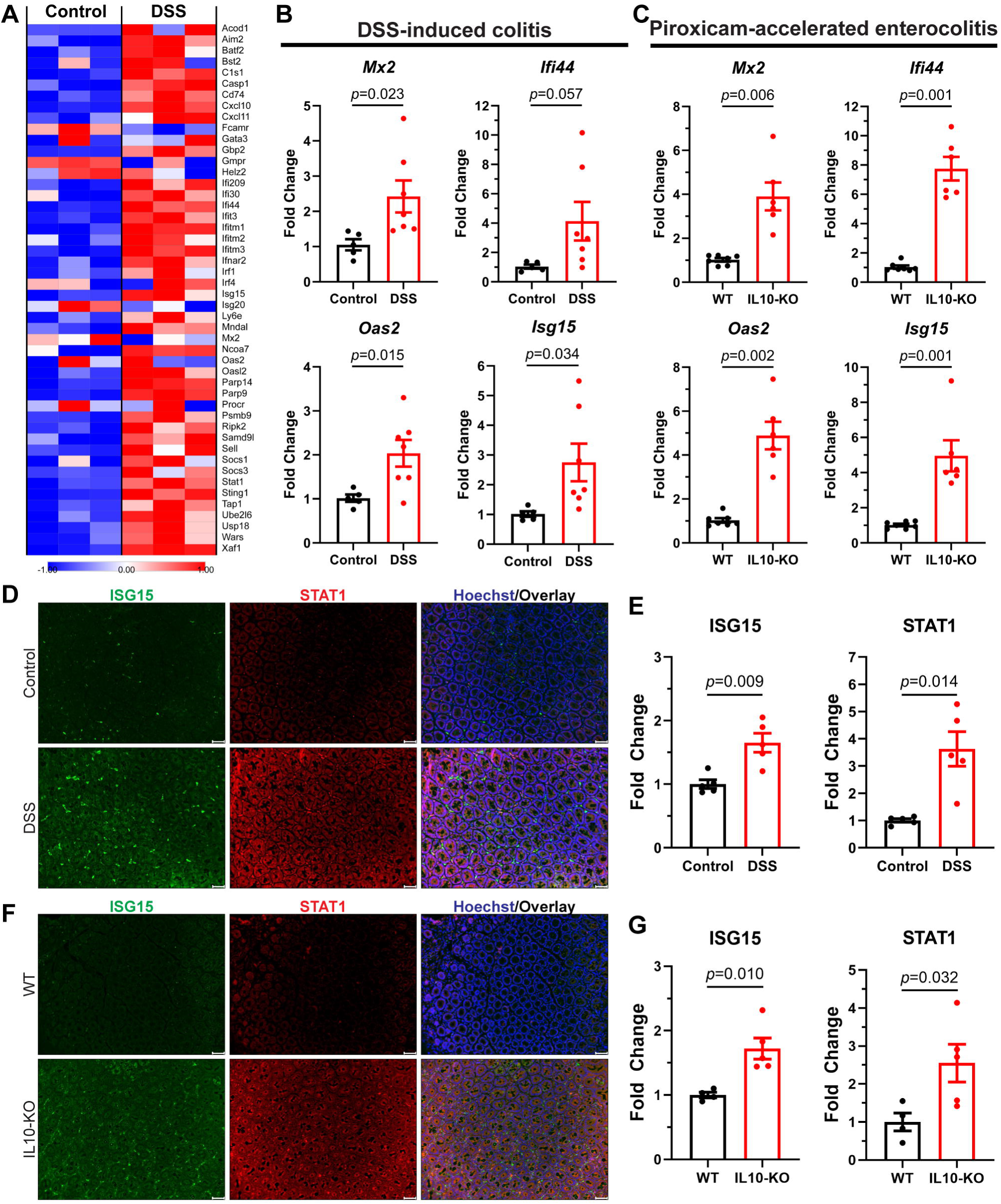
IFN-I signature genes are upregulated in murine models of colitis. **(A)** Heatmap showing the expression of 48 of the 54 IFN-I signature genes from Figure 2A in mouse colon with or without dextran sodium sulfate (DSS) exposure for 7 days, from re-analysis of a publicly available RNA-seq dataset (GSE131032). **(B-C)** qRT-PCR analysis of selected IFN-I signature genes from colonic tissues in DSS-induced colitis (2% DSS x 6 days followed by regular water x 3 days) in WT mice **(B)** and piroxicam-accelerated enterocolitis (piroxicam-fortified diet 200 ppm x 14 days) in IL10-KO mice **(C)**. Data were normalized to the mean of control samples (WT mice with no DSS exposure or WT mice receiving piroxicam-fortified diet, set as 1). n=5-6 per group. **(D-G)** Representative immunofluorescence images with quantification of ISG15 and STAT1 in DSS-induced colitis **(D-E)** and piroxicam-accelerated enterocolitis **(F-G)**. Immunofluorescence intensities were normalized to the mean of control samples or WT mice (set as 1). n=4-5 per group. Scale bar: 50 μm.

### Upregulated IFN-I signature genes are enriched in CD14^+^ myeloid cells

The colonic mucosa contains multiple cell types, including epithelial, immune, and stromal cells. While the bulk RNA-seq data integrates all signals in the tissue, it cannot ascribe the altered gene expression to a specific cell population, which is essential to understand potential pathogenic mechanisms. To resolve gene expression at the single cell level, we integrated publicly available single cell RNA sequencing (scRNA-seq) data from two human studies (GSE182270, GSE214695), which included tissue specimens from non-IBD healthy controls (n=9) and patients with active UC (n=12) (29, 30). UMAP representation of the transcriptomic data resolved the major cell populations expected to be present in the colonic mucosa **(Figure 4A-B).** Consistent with the clinical phenotype of UC, including epithelial ulceration and immune cell infiltration, the data demonstrated a decreased proportion of epithelial cells and corresponding increased proportion of immune cells in UC samples **(Figure 4C)**. We then performed pseudo-bulk RNA-seq analysis focusing our attention on the previously identified 54 IFN-I signature genes induced in active UC. Among all major cell clusters identified by our scRNA-seq analysis, CD14^+^ myeloid cells had the greatest number of significantly upregulated IFN-I signature genes in UC patients vs. healthy controls (11 genes with fold change >2 and FDR-adjusted *p* <0.05), followed by epithelial cells **(Figure 4D)**. This is illustrated by three representative genes, *ISG15*, *STAT1*, and *IFITM2*, which were notably increased in CD14^+^ myeloid cells from UC patients in our UMAP analysis **(Figure 4E)**.

**Figure 4.**
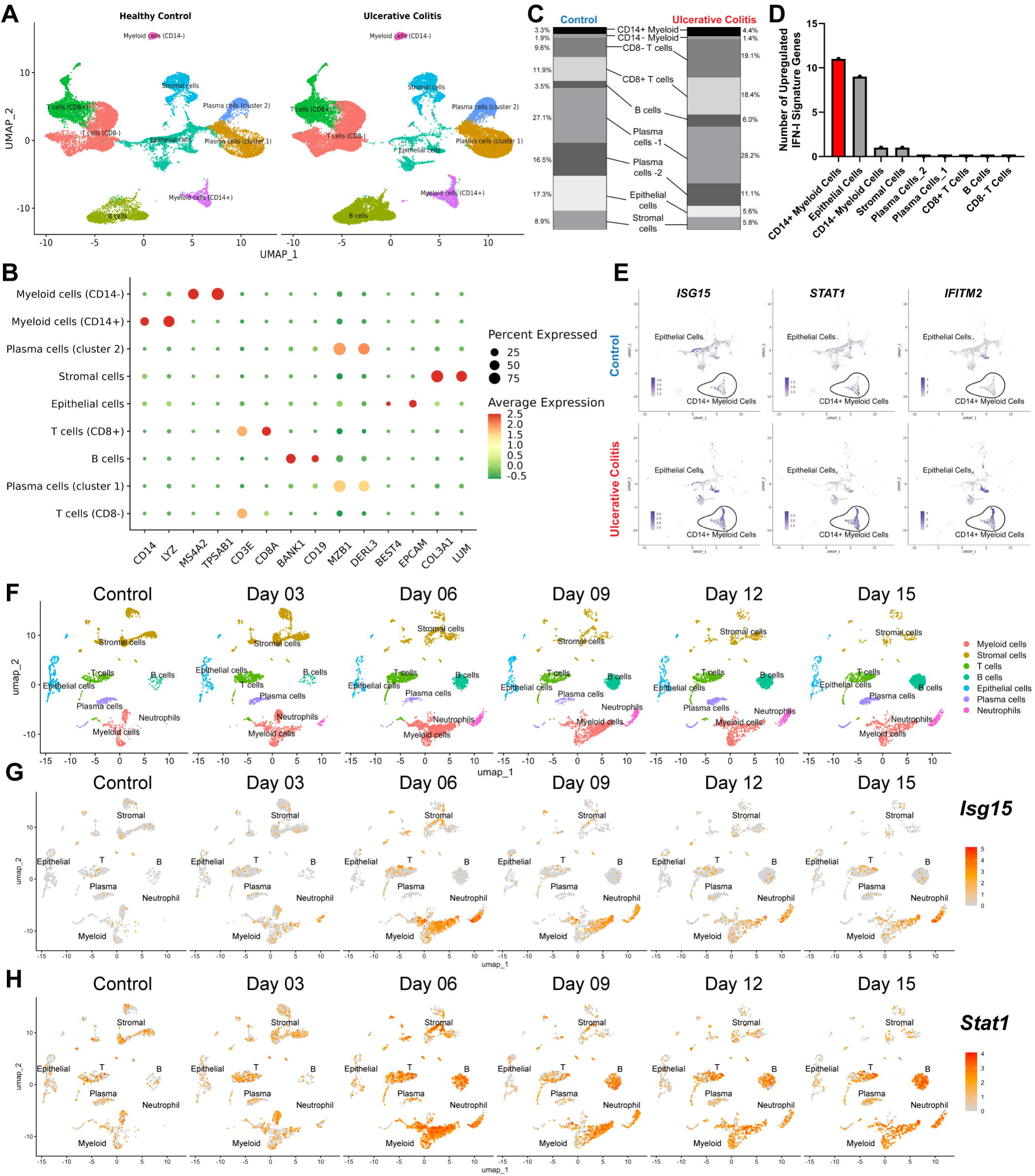
Upregulated IFN-I signature genes are enriched in CD14^+^ myeloid cells. **(A)** UMAP plots of scRNA-seq analysis of human colonic biopsies. Data from two publicly available scRNA-seq datasets (GSE182270, GSE214695) were integrated, including colonic biopsies from non-IBD healthy controls (n=9) and patients with active UC (n=12). **(B)** Dot plots of representative cluster markers in scRNA-seq analysis. **(C)** Stacked bar plots showing the proportion of each cell cluster in colonic biopsies from non-IBD healthy controls and patients with active UC. **(D)** Number of significantly upregulated IFN-I signature genes (fold change >2, FDR-adjusted *p* <0.05) among those identified in Figure 2A in each cell cluster. **(E)** Feature plots of representative IFN-I signature genes, *ISG15, STAT1,* and *IFITM2*, in epithelial cells and CD14^+^ myeloid cells. **(F)** UMAP plots of scRNA-seq analysis of DSS-induced colitis, from re-analysis of a publicly available scRNA-seq dataset (GSE148794) including mouse colon tissues collected at multiple time points (n=3 per time point) following 6 days of DSS and subsequent water recovery. **(G-H)** Feature plots of representative IFN-I signature genes, *Isg15* and *Stat1*, in all cell clusters at multiple time points.

To examine whether upregulation of IFN-I signature genes in murine colitis also occurs predominantly in myeloid cells, as seen in human IBD, we analyzed a publicly available scRNA-seq dataset (GSE148794) from a study of DSS-induced colitis (31). In this study, mice were exposed to DSS for 6 days and then switched to drinking water. DSS exposure was accompanied by a decrease in the epithelial and stromal compartments, with a progressive expansion in the immune cell compartment, particularly myeloid cells, including neutrophils (**Figure 4F**). Of note, *Isg15* and *Stat1* transcripts increased after DSS exposure and remained elevated after DSS withdrawal, predominantly in myeloid and other immune cell clusters **(Figure 4G-H)**.

### ISG-expressing myeloid cells express IFN-I and IFN-II receptors

As shown earlier, ISGs induced in active UC patients display significant overlap between IFN classes. To deconvolute potential contributions from IFN-I, IFN-II, and IFN-III, we examined the expression of IFN receptors in the scRNA-seq data, particularly in CD14⁺ myeloid cells and epithelial cells where upregulated ISGs were enriched. In human colonic biopsies, all IFN receptors were expressed in epithelial cells at low levels and without pronounced induction in active UC patients **(Figure 5A)**. On the other hand, CD14⁺ myeloid cells displayed substantial expression of *IFNAR1/IFNAR2* (IFN-I receptor) and *IFNGR1/IFNGR2* (IFN-II receptor), which increased in UC patient samples, but lacked *IFNLR1* expression (IFN-III receptor) **(Figure 5A)**. Next, we performed a similar analysis of IFN receptor expression in the murine model of DSS-induced colitis. *Ifnar1/Ifnar2* and *Ifngr1/Ifngr2* exhibited increased expression in immune cell clusters during and after DSS exposure, including myeloid cells and neutrophils, in agreement with the human data, whereas *Ifnrl1* expression was generally low and restricted to a small subset of cells **(Figure 5B)**. Altogether, these findings indicate that upregulated ISGs in myeloid cells that infiltrate inflamed tissues in human IBD and mouse colitis could be driven, at least in part, by IFN-I.

**Figure 5.**
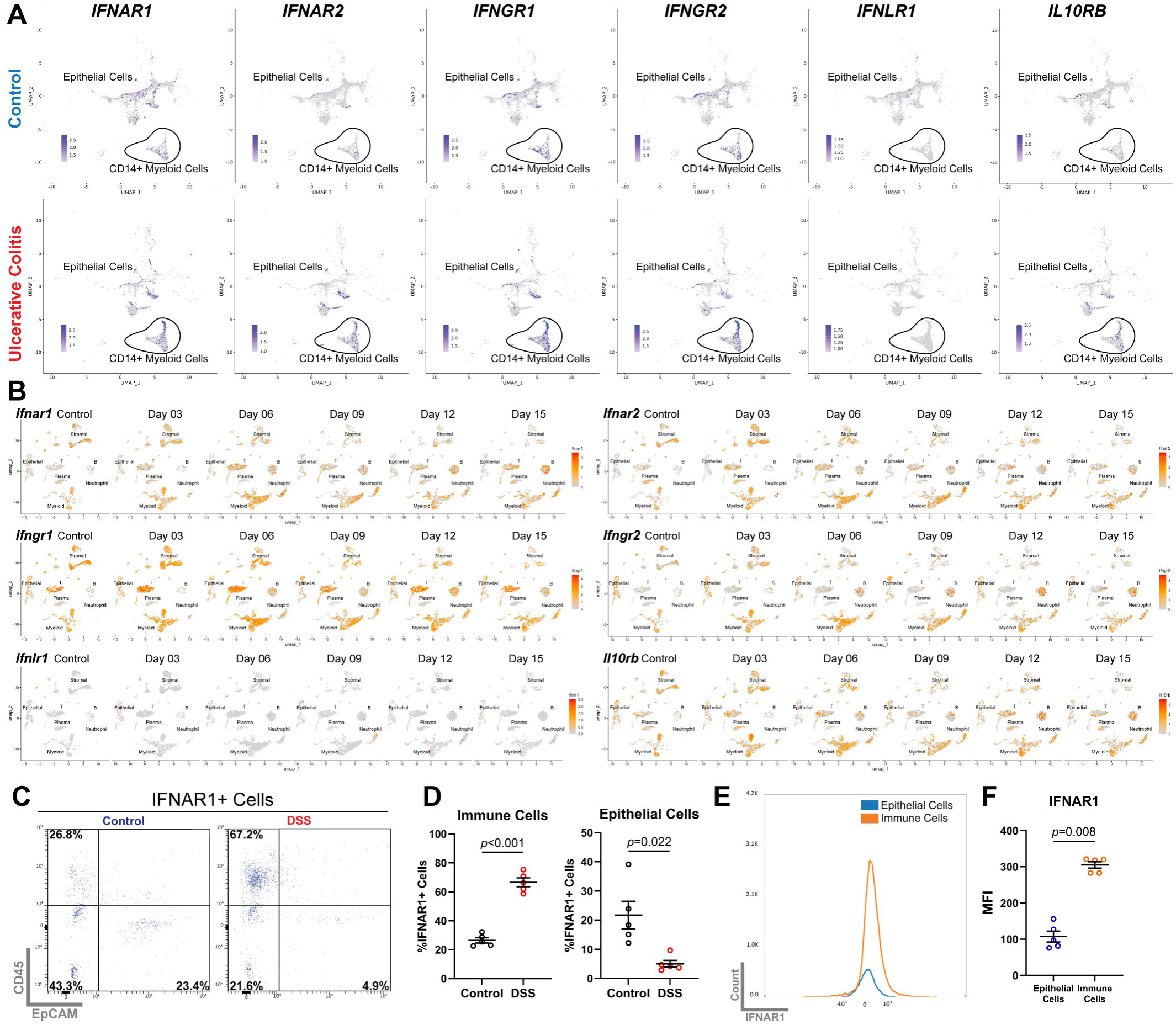
Expression of IFN receptors in human IBD and murine colitis. **(A)** Feature plots of IFN receptors, including IFN-I receptor (*IFNAR1/IFNAR2*), IFN-II receptor (*IFNGR1/IFNGR2*), IFN-III receptor (*IFNLR1/IL10RB*), in epithelial cells and CD14^+^ myeloid cells, from the scRNA-seq analysis of human colonic biopsies in Figure 4A. **(B)** Feature plots of IFN receptors, from the scRNA-seq analysis of DSS-induced colitis in Figure 4F. **(C)** Flow cytometry analysis of IFNAR1^+^ cells in mouse colon with or without DSS exposure, gated by immune cell marker CD45 and epithelial cell marker EpCAM. **(D)** Proportions of IFNAR1^+^ cells classified as immune cells (CD45^+^/EpCAM^-^) or epithelial cells (CD45^-^/EpCAM^+^) in mouse colon with or without DSS exposure. n=5 per group. **(E-F)** Flow cytometry analysis and quantification of median fluorescence intensity (MFI) of IFNAR1 expression in immune cells (CD45^+^/EpCAM^-^) and epithelial cells (CD45^-^/EpCAM^+^) from DSS-treated mouse colon.

At this point, we paid particular attention to IFN-I signaling and its receptor subunit, IFNAR1. We used flow cytometry to examine whether IFNAR1 in the colon showed differential expression following DSS-induced colitis. In uninflamed colon, only 26.8% of IFNAR1^+^ cells were immune cells (CD45^+^/EpCAM^-^), and the rest of IFNAR1^+^ population included stromal cells (CD45^-^/EpCAM^-^, 43.3%) and epithelial cells (CD45^-^/EpCAM^+^, 21.6%) **(Figure 5C-D)**. During DSS colitis, 67.2% of IFNAR1^+^ cells were immune cells **(Figure 5C-D)**, and only 21.6% and 4.9% were stromal cells and epithelial cells, respectively. Thus, consistent with the scRNA-seq data, our flow cytometry results indicated that the majority of IFNAR1^+^ cells in inflamed colonic tissues were immune cells. Importantly, in the context of colitis, immune cells also exhibited higher IFNAR1 median fluorescence intensity (MFI) than epithelial cells **(Figure 5E-F)**. Collectively, these results demonstrate an induction of IFN-I receptor along with upregulated IFN-I signature genes in the infiltrating immune cells in human IBD and murine colitis.

### Enhanced IFN-I signaling alters colonic immune homeostasis

While a pro-inflammatory role for IFN-II is well established, as outlined earlier, the potential role of IFN-I in mucosal inflammation remains unresolved with multiple conflicting results. To assess the contribution by excessive IFN-I signaling in immune homeostasis and experimental colitis, we resorted to a model of stabilizing expression of IFNAR1. IFNAR1 protein expression can be regulated through ubiquitination by the E3 ubiquitin ligase, SCF-βTrCP (3, 32). E3 recognition of IFNAR1 as a substrate is dependent on phosphorylation of the cytoplasmic tail of the receptor at serine 535 in the mouse protein **(Figure 6A)** (32). Thus, mice carrying a serine-to-alanine mutation at this site (IFNAR1^S535A^), hereafter referred to as SA mice, have been shown to exhibit defective IFNAR1 degradation via phosphorylation-dependent ubiquitination (33). In this study, we examined whether IFNAR1^S535A^ behaves as a gain-of-function (GOF) allele. Consistent with the predicted increased stability of the encoded mutant protein, surface expression of IFNAR1 was significantly higher in splenocytes from SA mice than WT littermates **(Supplemental Figure 2A-B)**. Moreover, mRNA expression of several IFN-I signature genes, including *Mx2, Ifi44, Oas2, and Isg15*, was significantly higher in spleen and colon tissues from SA mice compared to WT littermates **(Figure 6B)**. Similarly, bone marrow-derived macrophages (BMDMs) from SA mice exhibited higher mRNA levels of IFN-I signature genes at baseline, as well as following IFN-β or lipopolysaccharide (LPS) stimulation compared to WT BMDMs **(Figure 6C)**. These findings indicated that the SA model displays heightened basal tone and responsiveness to IFN-I stimulation, consistent with the higher IFNAR1 expression observed *in vivo*.

**Figure 6.**
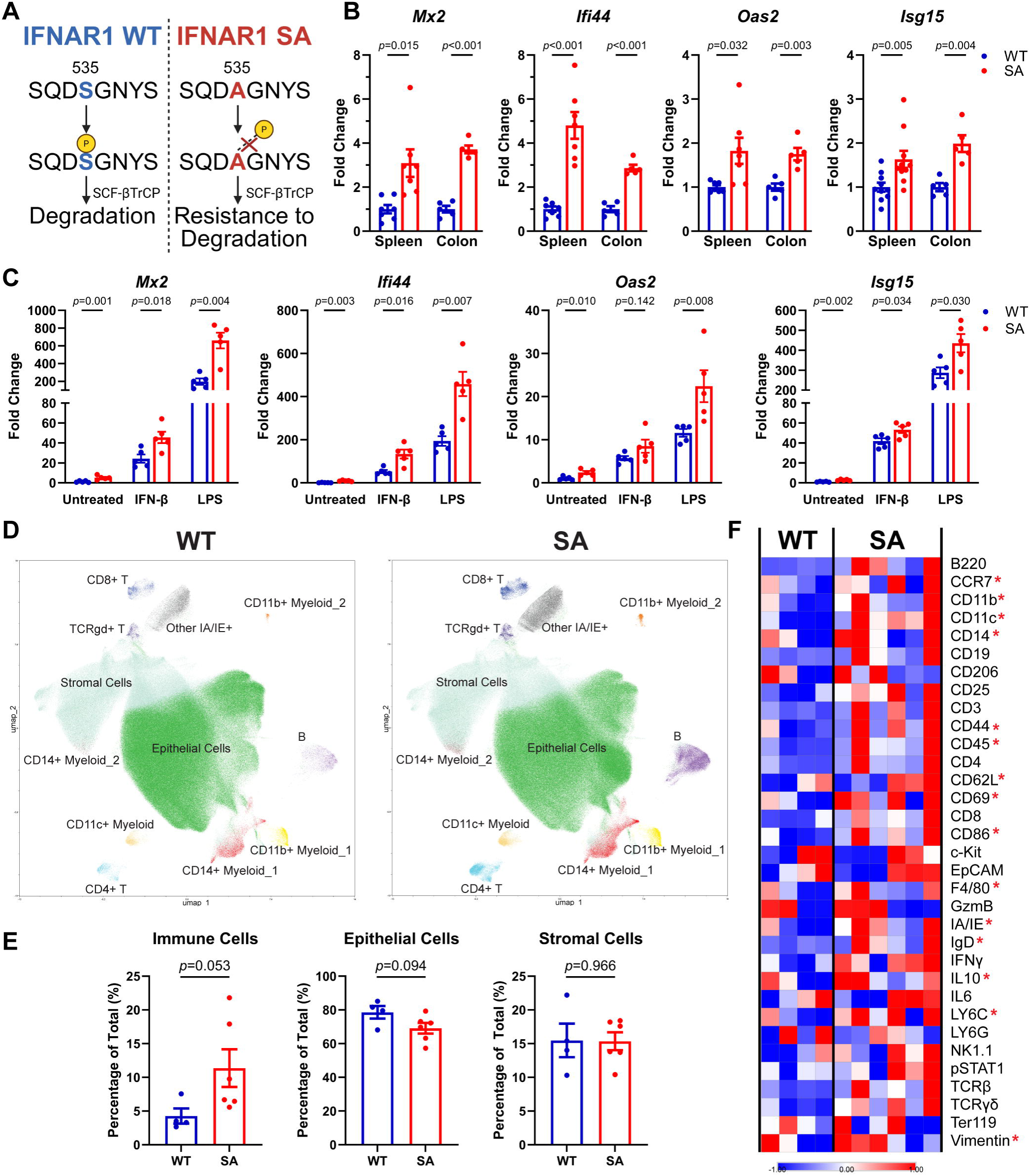
IFNAR1-SA mice exhibit elevated basal tone of IFN-I signaling and altered immune homeostasis. **(A)** Schematic illustrating how the serine-to-alanine substitution at position 535 of murine IFNAR1 protein affects phosphorylation and ubiquitination-dependent degradation. **(B)** qRT-PCR analysis of selected IFN-I signature genes in spleen and colon from WT and SA mice at baseline. Data were normalized to the mean of WT mice (set as 1). n=5-10 per group. **(C)** qRT-PCR analysis of selected IFN-I signature genes in bone marrow-derived macrophages (BMDMs) from WT and SA mice, at baseline (untreated) and after stimulation by interferon-β (IFNβ, 200 IU/mL x 8 hours) or lipopolysaccharides (LPS, 1 µg/mL x 8 hours). Data were normalized to the mean of WT BMDMs at baseline (set as 1). n=5-10 per group. **(D-E)** Mass cytometry (CyTOF) analysis of colon from WT (n=4) and SA (n=6) mice at baseline. **(D)** UMAP plots with cells colored by identity. **(E)** Proportions of immune cells (CD45^+^/EpCAM^-^), epithelial cells (CD45^-^/EpCAM^+^), and stromal cells (CD45^-^/EpCAM^-^). ns: not significant. **(F)** Heatmap showing the mean expression of target proteins in the antibody panel (Supplemental Table 4), with those having *p* <0.05 by SAM (Significance Analysis of Microarrays) for their median expression indicated by red asterisks.

To determine whether the elevated basal tone in SA mice is sufficient to induce intestinal inflammation, we examined mucosal cell composition using mass cytometry and an extensive antibody panel targeting 33 proteins **(Supplemental Figure 2C)**. We observed that immune cells (CD45^+^/EpCAM^-^) comprised a larger proportion of total cells in the colon of SA mice than WT mice, with a corresponding reduction in the proportion of epithelial cells (CD45⁻/EpCAM⁺), while stromal cells (CD45⁻/EpCAM⁻) showed no significant difference **(Figure 6D-E)**. These changes were also reflected in heatmap analysis, where levels of multiple immune cell markers were significantly higher in SA colon than in WT colon **(Figure 6F)**. Notably, CD86 and LY6C levels were significantly elevated in the colon from SA mice **(Figure 6F)**, indicating that in addition to their increased abundance, immune cells display a more activated pro-inflammatory state. A similar increase in CD86 and LY6C expression was also observed in the spleen. However, in this B-cell-dominant organ, the subcluster composition of immune cells was largely unchanged in SA mice **(Supplemental Figure 2D-F)**. Despite changes in cellular composition and increased expression of IFN-I signature genes, the gross architecture of the colon was unaffected at the histological level **(Supplemental Figure 2G)**. In addition, other than ISGs themselves (**Figure 6B**), baseline mRNA expression of inflammatory cytokines was not significantly different **(Supplemental Figure 2H)**, indicating a state of dysregulated immune homeostasis without clinical colitis in SA mice.

### SA mice are more susceptible to experimental colitis

While SA mice do not exhibit a spontaneous colitis, we asked whether they might display increased susceptibility to chemically induced colitis, similar to their heightened inflammatory response in other experimental settings (33, 34). First, we assessed the response of SA mice to DSS-induced colitis. In this model, SA mice developed more severe disease, showing greater weight loss, higher disease activity scores, more prominent colon shortening, and elevated transcript levels of inflammatory cytokines (*Il1b*, *Il6)* and IFN-I signature genes (*Mx2*, *Pyhin1*) **(Figure 7A–D)**. Next, we assessed the contribution of excessive IFN-I signaling on the development of enterocolitis in IL10-KO mice. IL10-KO mice are known to develop spontaneous enterocolitis (35), consistent with severe early-onset IBD seen in humans with loss-of-function (LOF) mutations in *IL10* or the genes encoding its receptor (36). Depending on the underlying mouse strain and vivarium conditions, the phenotype may have variable penetrance and take many weeks to develop (35). For this reason, we used a piroxicam-accelerated model, where IL10-KO mice were exposed to piroxicam-fortified diet, which results in 100% penetrance in the span of 7-21 days (28). Through intercrossing, we generated IL10-KO mice with the *Ifnar1^S535A^* allele and evaluated their susceptibility to piroxicam-accelerated enterocolitis compared to animals with the WT *Ifnar1* allele. Using the same assessment methods as above, we observed more profound weight loss and more severe disease activity in SA mice compared to littermate WT controls, with colon length trending in the same direction **(Figure 7E–G).** In addition, transcript levels of inflammatory cytokines and IFN-I signature genes were more elevated in the colon of SA mice than WT littermates. Altogether, these results demonstrated increased susceptibility to experimental colitis in the context of excessive IFNAR1-mediated signaling.

**Figure 7.**
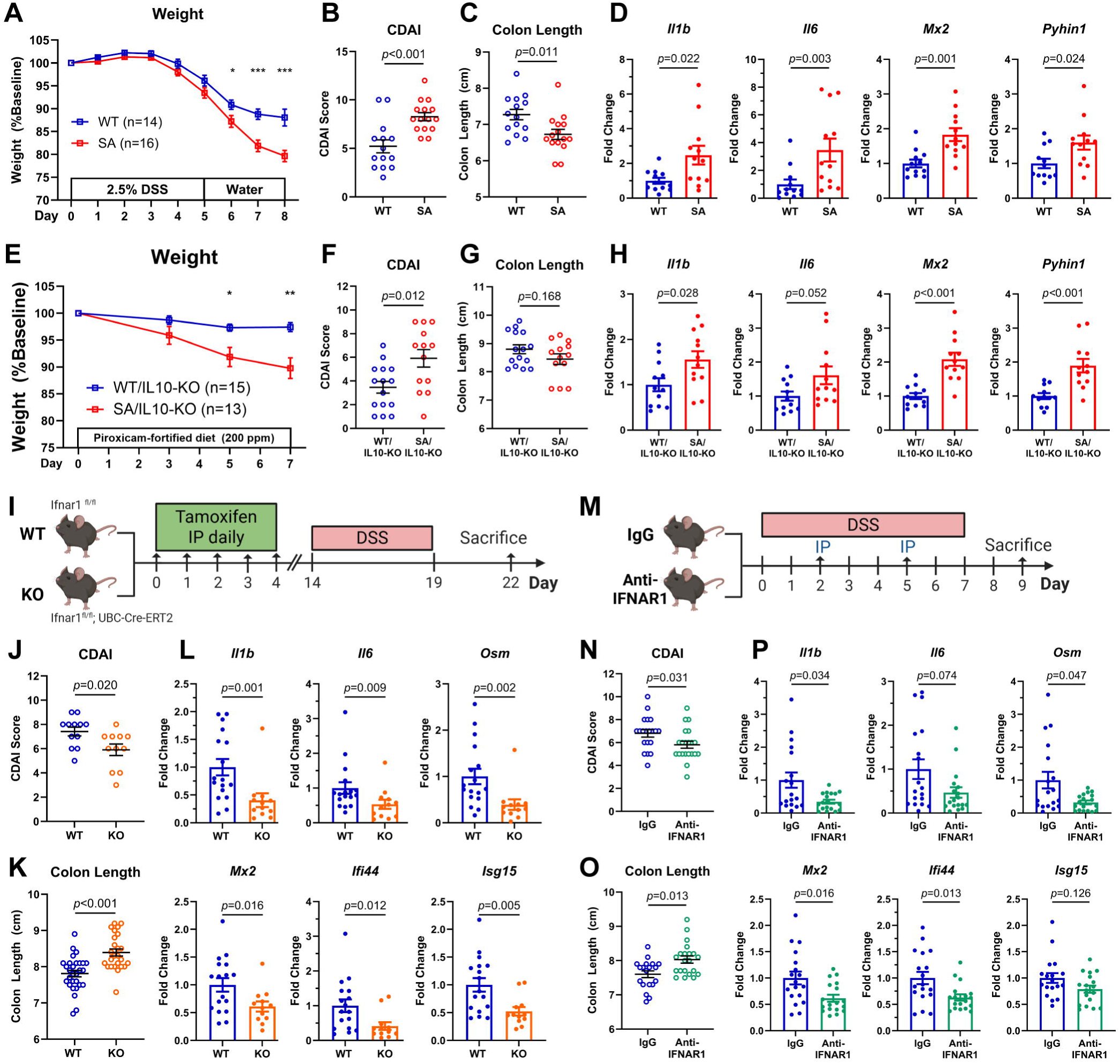
IFNAR1-SA mice are more susceptible to experimental colitis, while post-natal IFNAR1 inhibition is protective (A-H) Weight change **(A and E)**, clinical disease activity index (CDAI) **(B and F)**, colon length **(C and G)**, and qRT-PCR analysis of selected inflammatory cytokines and IFN-I signature genes **(D and H)** in colon in DSS-induced colitis **(A-D)** and piroxicam-accelerated enterocolitis **(E-H)**. **(I-P)** Schematics of experimental design **(I and M)**, clinical disease activity index (CDAI) **(J and N)**, colon length **(K and O)**, and qRT-PCR analysis of selected inflammatory cytokines and IFN-I signature genes **(L and P)** to assess the effect of tamoxifen-induced knockout of IFNAR1 **(I-L)** and administration of blocking anti-IFNAR1 antibody **(M-P)** on DSS-induced colitis.

### Post-natal inhibition of IFN-I signaling protects against experimental colitis

IFN-I signaling has been shown to regulate multiple aspects of immune cell development, differentiation, recruitment, and activation (37, 38). Thus, the SA model could reflect a change in immune development that ultimately lowers the threshold for intestinal inflammation. To assess whether IFNAR1-dependent signaling can exert an effect on intestinal inflammation through non-developmental mechanisms, we asked whether post-natal attenuation of IFN-I signaling protects against experimental colitis. First, we used a transgenic mouse model of tamoxifen-inducible *Ifnar1* deletion. In this model, mice carrying a floxed *Ifnar1* allele were mated with the *Ubc^CreERT2^* transgenic line, which expresses Cre recombinase in all tissues but can only mediate effective gene deletion upon treatment with tamoxifen **(Figure 7I)**. Unlike germline IFNAR1-KO mice, these conditional IFNAR1-KO mice maintain normal IFN-I signaling during development, avoiding potential effects of IFN-I loss on cell differentiation. Following tamoxifen administration, the surface expression of IFNAR1 in splenocytes, measured by flow cytometry, was significantly reduced **(Supplemental Figure 3A-B)**. In addition, qRT-PCR confirmed near complete loss of *Ifnar1* mRNA in the colon tissue **(Supplemental Figure 3C)**, confirming the validity of the model. Next, we assessed the susceptibility of these animals to DSS-induced colitis. We observed that post-natal global deletion of *Ifnar1* conferred protection against DSS-induced colitis, reflected predominantly by reduced disease activity score, preserved colon length, and reduced colonic mRNA expression of inflammatory cytokines and IFN-I signature genes **(Figure 7J-L)**. Curiously, weight reduction during colitis was nearly identical between groups **(Supplemental Figure 3D)**. Next, we evaluated whether pharmacologic inhibition of IFN-I signaling could also ameliorate DSS-induced colitis. To this end, we used a blocking antibody against mouse IFNAR1. Of note, the corresponding anti-human IFNAR1 antibody has been approved for the treatment of SLE (36). WT mice received intraperitoneal injections of the anti-IFNAR1 antibody or isotype control IgG during DSS exposure **(Figure 7M)**. Similar to post-natal deletion of *Ifnar1*, the group that received the blocking antibody exhibited lower disease activity index, less colon shortening, and lower levels of inflammatory cytokines and IFN-I signature genes **(Figure 7N-P)**, consistent with a therapeutic effect of IFNAR1 blockade against DSS-induced colitis. However, the effect of the blocking antibody on body weight was minimal and not significant **(Supplemental Figure 3E)**, suggesting that systemic responses to disease activity, such as anorexia and weight loss, may not be mediated by IFN-I signaling. Altogether, both transgenic and pharmacologic models of acute IFNAR1 inhibition produced highly concordant phenotypes consistent with a pathogenic role of IFN-I signaling in experimental colitis.

## DISCUSSION

While IFN-I signaling has been extensively studied in autoimmune and infectious diseases, its role in IBD has remained incompletely defined, with a prevailing view in the field that it may be protective due to its immunomodulatory and anti-viral functions (7, 23, 39). Using genetic and pharmacologic approaches to modulate IFNAR1-dependent signaling, our work demonstrates that increased IFN-I signaling promotes intestinal inflammation whereas acute inhibition of IFN-I signaling attenuates experimental colitis, in agreement with a number of recent studies (40, 41). We speculate that several mechanisms may mediate the pathogenic effects of IFN-I signaling in IBD, including immune cell infiltration (37, 42, 43) and immune cell differentiation and activation (37, 44), both of which were observed in the tissues of SA mice. In addition, persistent IFN-I signaling may interfere with epithelial cell differentiation and promote apoptosis (45), which may have limited impact during the acute phase of colitis but could impair tissue regeneration in chronic inflammation.

Our pro-inflammatory view of IFN-I is consistent with genetic evidence in human IBD. First, genetic studies have found that variants in several genes in the IFN-I pathway, such as *JAK1*, *TYK2*, *IFNAR1*, *STAT1,* are associated with increased risk of IBD (46, 47). In addition, examples of monogenic immune disorders with intestinal inflammation also implicate IFN-I activation in disease pathogenesis. For instance, X-linked reticulate pigmentary disorder (XLPDR), a rare genetic disorder where more than 70% of patients develop infantile-onset ulcerative colitis, is characterized by activation of IFN-I signaling due to a hypomorphic mutation in the *POLA1* gene (18). Moreover, targeting of JAK1, a key kinase in the IFN-I signaling pathway, was effective in controlling intestinal inflammation in a patient with XLPDR, further supporting a pathogenic contribution of IFN-I in the disorder (48).

Recognition of a potential pathogenic role of IFN-I in IBD provides a new perspective on disease mechanisms and has important clinical implications for therapeutic targeting. First, while a significant ISG signature in IBD biopsies and blood samples has been previously reported, it was largely understood as a bystander response to mucosal inflammation and its clinical significance was not fully elucidated (49–52). Our data indicate that this ISG signature serves as a marker of disease activity, given higher levels observed in colonic biopsies from active UC patients compared with inactive UC patients and non-IBD controls. Beyond that, an ISG signature may also reflect resistance to certain types of treatment, as it has been reported that patients refractory to anti-TNF therapy have more pronounced ISG expression than responsive patients prior to receiving therapy (49, 50). This suggests that in certain IBD patients, a prominent ISG signature may reflect a disease process driven primarily by IFN signaling that may be less responsive to anti-TNF therapy. In view of our results, it stands to reason that such patients may be potentially responsive to IFN-inhibitory therapies such as JAK1 inhibitor, which has already been approved for IBD treatment (53, 54), as well as TYK2 inhibitor, which is currently being evaluated for this indication in clinical trials (55). In addition, it is also important to consider that the therapeutic benefit from JAK1 inhibitor may be derived from inhibition of IFNAR1-dependent signaling, and therefore, it is also intriguing to hypothesize that patients displaying elevated ISG expression may experience a more significant response to this treatment modality. Lastly, our data support the potential therapeutic utility of anti-IFNAR1 antibody in IBD. This therapy has been approved for treating patients with SLE (36), and its therapeutic efficacy in IBD patients merits clinical testing in the future.

Our experimental observations in mouse models stand in contrast with a few studies in the literature, where inconsistent results in different contexts have been reported. For example, our data in mice that underwent post-natal deletion of *Ifnar1* are the opposite of the phenotypes reported for germline *Ifnar1* deletion, which results in variable colitis sensitivity rather than disease protection (20–22). While the phenotype observed in the germline knockout models were largely attributed to the loss of the protective effects of IFN-I, our results suggest that this may also reflect developmental defects due to lifelong deprivation of IFN-I signaling. That said, others have reported that systematic IFN-β administration in mice can ameliorate DSS-induced colitis (20); however, the reported results were largely focused on disease activity and did not include molecular surrogate markers of inflammatory activity. Moreover, IFN-β use in humans demonstrated no beneficial activity (13). Some studies have also reported opposing effects of IFNAR1 global deficiency depending on the DSS dose used (21), and still others have demonstrated that concurrent deficiency of IFN-I and IFN-III receptors in mice results in severe colitis due to poor epithelial restitution (22). Again, because of the potential developmental defects secondary to loss of IFN-I signaling, we postulate that acute manipulation of IFN-I signaling is likely to provide a more precise assessment of its function during intestinal inflammation.

In summary, our findings reveal a pathogenic role of IFN-I signaling in IBD, providing new insight into its complex function in intestinal homeostasis. Further studies are needed to delineate whether IFN-I signaling coordinates with other inflammatory pathways and to identify the specific ISGs and cell types responsible for this pathogenic effects, which will be critical for advancing our understanding of IBD pathogenesis and informing the development of novel therapeutic strategies.

## METHODS

### Sex As A Biological Variable

Human studies included participants of both sexes, and sex was not considered a key biological variable. Animal studies included both male and female mice. In DSS-induced colitis experiments, male mice were used to minimize biological variability related to sex and to align with the existing literature in which the model of DSS-induced colitis has been predominantly studied using male mice (56). For similar reasons, in piroxicam-accelerated enterocolitis experiments, female mice were used because prior studies used female mice (28, 57).

### Study Approval

The study protocol for human studies was in accordance with the ethical guidelines established in the Declaration of Helsinki and was approved by the Institutional Review Board (IRB). All participants were recruited at the University of Texas Southwestern Medical Center at Dallas, TX, and written informed consents were obtained. For animal studies, all mice were treated in compliance with the protocol approved by the Institutional Animal Care and Use Committee (IACUC).

### Reagents

Antibodies used in this study are listed in **Supplementary Table 3 and 4**. Other reagents are described in the relevant Methods sections, with additional details available upon request.

### Human Subjects

Patient specimens were obtained from the University of Texas Southwestern Medical Center (UT Southwestern) biorepository for the study of inflammatory GI disorders (IRB number STU112010-130). For the bulk RNA-seq study, the biorepository database was queried to identify sigmoid colon biopsies obtained from patients with UC during diagnostic colonoscopy and from non-IBD controls during screening colonoscopy. Disease activity was determined using endoscopic findings and pathology reports, with active disease defined as an endoscopic Mayo score of 2-3 with histologic evidence of active colitis, and inactive disease defined as an endoscopic Mayo score of 0 with histologic evidence of inactive colitis. For the immunofluorescence study, rectal biopsies were prospectively collected from patients with acute severe UC undergoing sigmoidoscopy or colonoscopy, as well as from non-IBD healthy controls undergoing screening colonoscopy.

### Mice

We used >8 weeks old, age- and gender-matched littermate mice with similar baseline body weights for all experiments. WT mice (strain #000664), SA mice (*Ifnar1*^S526A^ knock-in, strain #035564), IL10-KO mice (strain #002251), *Ifnar1*^fl/fl^ mice (strain #028256), *Ubc*-CreERT2 mice (strain #007001) were purchased from the Jackson Laboratory and maintained in the animal facility at UT Southwestern. All strains came in the C57BL/6J genetic background. Mice were maintained by heterozygous breeding. SA mice were compared with their WT littermates, and *Ifnar1*^fl/fl^ mice were compared with their littermates with *Ubc*-CreERT2. Genotypes of all mice were confirmed both before and after each experiment. Primer sequences used for genotyping are listed in **Supplemental Table 5**.

### Murine Models of Colitis and in vivo Studies

Unless otherwise specified, for DSS-induced colitis, mice were given 2.5% (w/v) dextran sodium sulfate (Thermo Fisher Scientific, J63606.22, molecular weight 35,000-50,000) in drinking water for 5-7 days, followed by regular drinking water for 2-3 days. For piroxicam-accelerated enterocolitis, mice were given a piroxicam-fortified diet (200 ppm, Sigma-Aldrich, P0847, prepared by Inotiv) for 7-14 days. Mice had *ad libitum* access to DSS-containing water or piroxicam-fortified diet during the respective treatment periods, but did not have access to regular water or standard chow while on these treatments.

The scoring system for calculating clinical disease activity index (CDAI) was adapted from Wirtz, *et al* (58) and is detailed in **Supplemental Table 6**. The total CDAI score was calculated by adding individual scores from weight loss, stool consistency, and presence of blood in stool. Fecal occult blood was assessed using the Hemoccult Sensa Single Slides System (Beckman Coulter).

For tamoxifen-induced gene knockout, *Ifnar1*^fl/fl^ mice with or without *Ubc*-CreERT2 received intraperitoneal injections of 100 μL tamoxifen (20 mg/mL in corn oil, Sigma-Aldrich, T5648) daily for 5 consecutive days and were subject to experiments 10-12 days later. For pharmacologic inhibition of IFNAR1, mice were intraperitoneally injected with 0.5 mg anti-mouse IFNAR1 antibody (BioXcell, BE0241) or mouse IgG1 isotype control (BioXcell, BE0083) on day 2 and day 5 after DSS exposure.

### Bone Marrow-Derived Macrophages (BMDMs)

Bone marrow cells were harvested from mouse leg bones and plated in culture media consisting of 45% DMEM, 20% FBS, and 35% L929 conditioned media (prepared by collecting media from confluent L929 cells cultured in 90% DMEM, 10% FBS, and 1 g/dL GlutaMax, at a ratio of 2.8 mL of media per square centimeter of area of confluent L929 cells). This culture condition supported differentiation of bone marrow cells into BMDMs. Media were refreshed every 2-3 days, and BMDMs were used for experiments 10 days after plating. For stimulation studies, BMDMs were treated with interferon-β (IFN-β, 200 IU/mL x 8 hours) or lipopolysaccharides (LPS, 1 µg/mL x 8 hours).

### RNA Isolation and Quantitative Reverse Transcription PCR (qRT-PCR)

Samples were stored in RNAlater Stabilization Solution (Thermo Fisher Scientific, AM7021) at -20 °C. Tissues were homogenized using a Spex SamplePrep 1600 MiniG tissue homogenizer. Total RNA was extracted using the RNeasy Mini Kit (Qiagen) according to the manufacturer’s protocol. Complementary DNA was synthesized using SuperScript VILO Master Mix (Thermo Fisher Scientific). qRT-PCR was performed using PowerTrack SYBR Green Master Mix (Thermo Fisher Scientific) on a QuantStudio 7 Pro real-time PCR system (Thermo Fisher Scientific). We used 5 ng RNA-equivalent complementary DNA for each reaction, and each sample was run in triplicate. Primers used for qRT-PCR studies are listed in **Supplemental Table 5**. The relative fold changes in mRNA levels were determined using the 2 ^-ΔΔCT^ method with human *GAPDH* or mouse *Gapdh* simultaneously studied and used for normalization as the internal control.

### Bulk RNA Sequencing and Analysis

Bulk RNA sequencing was performed by Genewiz (Azenta Life Sciences). In brief, total RNA was isolated from colonic biopsies as described above, and RNA integrity was assessed using a TapeStation (Agilent Technologies). RNA sequencing libraries were prepared using the NEBNext Ultra II RNA Library Prep for Illumina. Samples meeting quality control criteria were used for library preparation using a poly(A)-selected, stranded mRNA protocol, followed by high-throughput sequencing on an Illumina NovaSeq instrument according to manufacturer’s instructions. Sequencing data were processed using a custom analysis pipeline developed by the UT Southwestern BioHPC facility. Reads were aligned to the reference genome (GRCh38 for human) and gene-level counts were generated. Differential expression analysis was performed using DESeq2, with a fold-change cutoff of >2 and FDR-adjusted *p* <0.05.

### Single Cell RNA Sequencing Analysis

Publicly available single-cell RNA sequencing data were obtained from the Gene Expression Omnibus (GEO) and analyzed in R using the Seurat package (Seurat v4.3.0 in R v4.2.2 for human datasets and Seurat v5.3.1 in R 4.5.1 for mouse datasets). Cells were filtered using standard quality control criteria (200-4,000 detected genes with <25% mitochondrial gene expression for human datasets; 200-6,000 detected genes with <15% mitochondrial gene expression for mouse datasets). Datasets were normalized and integrated using Seurat’s integration workflow, followed by principal component analysis. The top 30 principal components were used for downstream analysis of human samples and the top 20 for mouse samples. Clustering was performed using a resolution of 0.1 for human data and 0.03 for mouse data. Cluster identities were assigned based on the top 20 differentially expressed genes identified for each cluster using the FindMarkers function, together with established canonical cell-type markers. Pseudobulk differential expression analysis was performed for each cluster using DESeq2. Differentially expressed genes were defined as those with a fold change >2 and an FDR-adjusted *p* value <0.05.

### Histology and Immunofluorescence

Slides were prepared from formalin-fixed, paraffin-embedded tissue blocks and stained with hematoxylin and eosin (H&E) using a standard protocol. For immunofluorescence, slides were warmed at 60°C for 20 minutes and deparaffinized using the sequential xylene method, followed by rehydration with decreasing alcohol concentrations and heat-induced antigen retrieval in citrate solution (Sigma-Aldrich, C9999). Then, slides were incubated with blocking buffer (3.5% goat serum in PBS) at room temperature for 30 minutes, followed by primary antibodies diluted in blocking buffer at 4 °C overnight **(Supplemental Table 3)**. The next day, slides were incubated in blocking buffer at room temperature for 30 minutes, secondary antibodies diluted in blocking buffer at 4 °C for 60 minutes, and Hoechst 33258 (Sigma Aldrich, 94403) for 20 minutes **(Supplemental Table 3)**. Finally, slides were mounted with SlowFade Gold Antifade Reagent (ThermoFisher Scientific, S36937).

Images were acquired using a Keyence BZ-X810 fluorescence microscope. Immunofluorescence intensity was quantified using Fiji (version 2.16.0), and intensities of proteins of interest in each image were normalized to nuclear counts determined by Hoechst 33258 staining. The mean value of 2-3 biopsies per human participant or 3 colon areas per mouse was used to represent a biological replicate. For comparisons between groups, the mean values of non-IBD controls, control mice without DSS exposure, or WT mice after piroxicam exposure were used as the reference (set as 1).

### Single Cell Isolation

Single cell suspensions were prepared from mouse spleen or colon tissues. For spleen, samples were gently dissociated through a 100 μm cell strainer and incubated with RBC lysis buffer (Thermo Fisher Scientific, 00-4333-57) at room temperature for 5 minutes. For colon, samples were opened longitudinally, rinsed with PBS, washed with HBSS supplemented with 1 mM EDTA (Thermo Fisher Scientific, J15694.AE), 2 mM HEPES (Thermo Fisher Scientific, 15630080), and 1 mM DTT (Sigma Aldrich, D0632) at 37 °C for 20 minutes, chopped into small pieces, and then digested in RPMI 1640 media containing 0.25 mg/mL type III collagenase (Sigma-Aldrich, C2139) and 2 units/mL DNase I (Sigma-Aldrich, D4527) at 37 °C for 1 hour with agitation. Digested cell suspensions were filtered through a 100 μm cell strainer and centrifuged in 40% Percoll. Finally, cells were resuspended in 0.5% BSA in PBS for subsequent studies.

### Mass and Flow Cytometry

For flow cytometry, cells were resuspended in blocking buffer containing 0.5% BSA and CD16/CD32 antibody (Thermo Fisher Scientific, 14-0161-81) to block Fc receptors. Then, cells were stained with fluorophore-conjugated antibodies **(Supplemental Table 4)** and viability dye (LIVE/DEAD™ Fixable Yellow Dead Cell Stain, Thermo Fisher Scientific, L34967) at 4°C for 20 minutes in dark, fixed using BD Cytofix/Cytoperm (BD Biosciences, 554714) at 4°C for 20 minutes in dark, and resuspended in 0.5% BSA in PBS.

For mass cytometry, cells were first incubated at room temperature with Cell-ID Cisplatin (Standard BioTools) for 20 minutes and then with TruStain FcX (BioLegend) for 10 minutes. Then, cells were incubated with metal-conjugated primary antibodies that target surface proteins **(Supplemental Table 4)** at room temperature for 30 minutes, followed by fixation and permeabilization using Maxpar Fix I buffer for 20 minutes and MaxPar Perm-S buffer (Standard BioTools) for 30 minutes. Intracellular proteins were subsequently stained with metal-conjugated antibodies at room temperature for 30 minutes **(Supplemental Table 4)**. Cells were fixed with 1.6% formaldehyde for 10 minutes, incubated with Cell-ID Intercalator-Ir (Standard BioTools) at 4°C overnight, and resuspended in Maxpar Cell Acquisition Solution (Standard BioTools).

Flow cytometry data were acquired on an Aurora flow cytometer (Cytek Biosciences) using SpectroFlo software (version 3.3.0) after calibration with Submicron Bead Calibration Kit (Bangs Laboratories). Mass cytometry data were acquired on a Helios mass cytometer (Standard BioTools). Data were analyzed using Omiq.

### Statistics

Data are shown as mean ± SEM, unless otherwise specified. Statistical analyses were conducted using GraphPad Prism (version 10.6.1). Depending on data distribution assessed by the Shapiro–Wilk test, statistical analysis for two-group comparisons were performed using the Welch’s t-test or the Mann–Whitney test, and for multiple-group comparisons using the one-way ANOVA or the Kruskal–Wallis test, unless otherwise indicated. A *p* value less than 0.05 was considered significant.

## Data Availability

The publicly available RNA-seq and scRNA-seq data re-analyzed in this study are available from the GEO database under accession numbers GSE182270, GSE214695, GSE131032, GSE148794. The bulk RNA-seq data for Figure 1 in raw FASTQ format are deposited at GEO repository. Additional data are available upon reasonable request.

## AUTHOR CONTRIBUTIONS

JY and EB conceived the project, designed the experiments, interpreted the results, and wrote the manuscript. JY and RGP performed experiments. LSD and ET revised the manuscript and provided critical intellectual input. EB supervised the study.

## ACKNOWLEDGMENTS

We thank the Flow Cytometry Core, the Tissue Management Shared Resource, and the Animal Resource Center (ARC) at UT Southwestern Medical Center for technical support. This research was supported in part by the computational resources provided by the BioHPC supercomputing facility located in the Lyda Hill Department of Bioinformatics, UT Southwestern Medical Center. We thank all participants in the biorepository, and the research coordinators, fellows, and faculty members in the Division of Digestive and Liver Diseases at UT Southwestern Medical Center for their assistance with specimen collection,

---

**Supplemental Figure 1.**
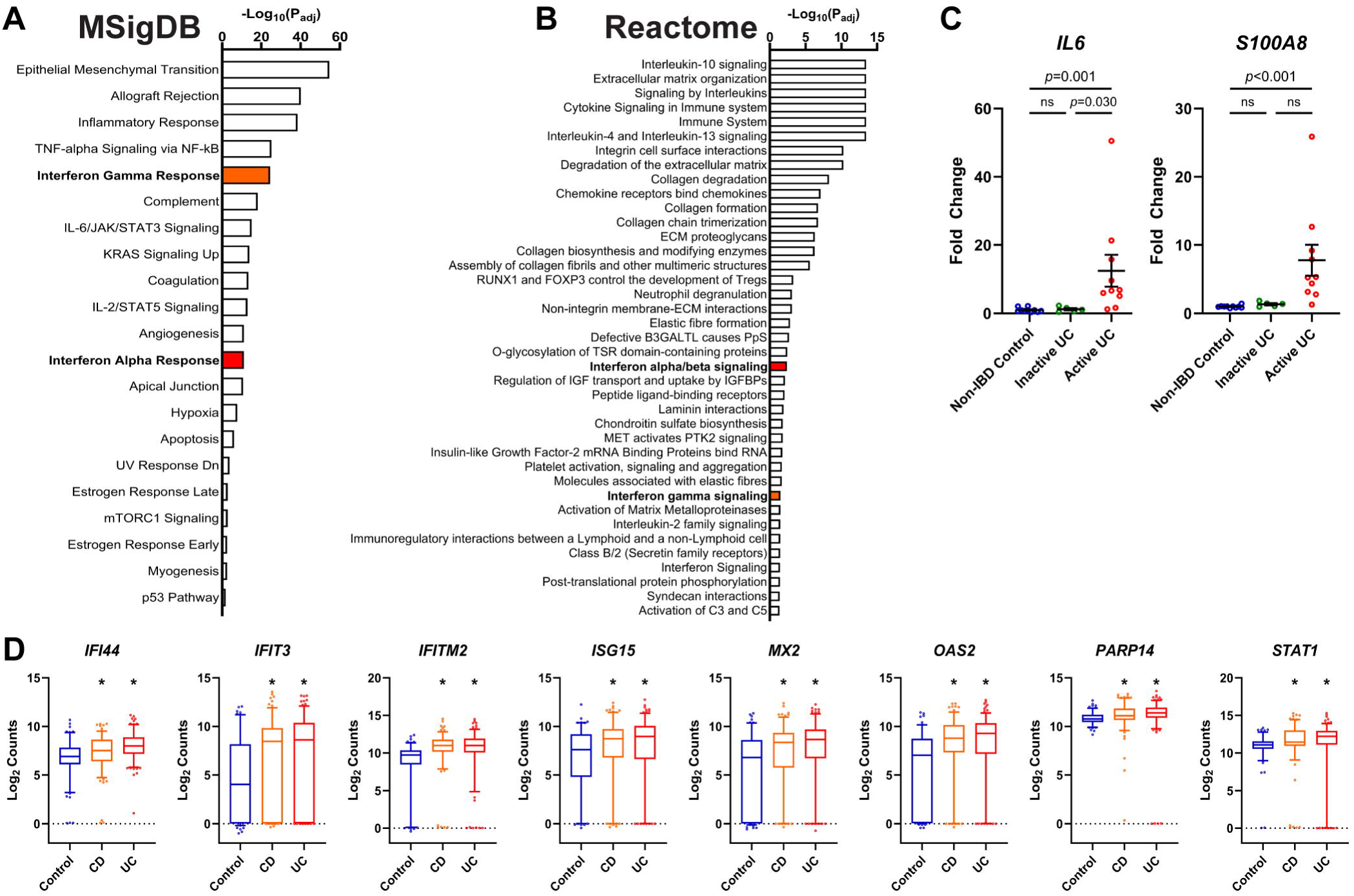
IFN-I signaling is activated in colonic biopsies from patients with inflammatory bowel disease. (A-B) Pathway enrichment analysis of the upregulated DEGs identified in Figure 1A using Molecular Signatures Database (MSigDB) **(A)** and Reactome Pathway Database **(B)**. **(C)** qRT-PCR validation of selected inflammatory cytokines (*IL6*, *S100A8*) in colonic biopsies. n=6-11 per group. ns: not significant. **(D)** Analysis of selected IFN-I signature genes (*IFI44*, *IFIT3*, *IFITM2*, *ISG15*, *MX2*, *OAS2*, *PARP14*, *STAT1*) in the IBD TaMMA database comparing colonic samples from non-IBD controls (n=123), patients with Crohn’s disease (CD, n=156), and patients with ulcerative colitis (UC, n=169). Log_2_-transformed adjusted counts are presented as box-and-whisker plots (center line indicates the median, whiskers represent 5^th^-95^th^ percentile). Asterisks indicate FDR-adjusted *p* values <0.05 compared with non-IBD controls.

**Supplemental Figure 2.**
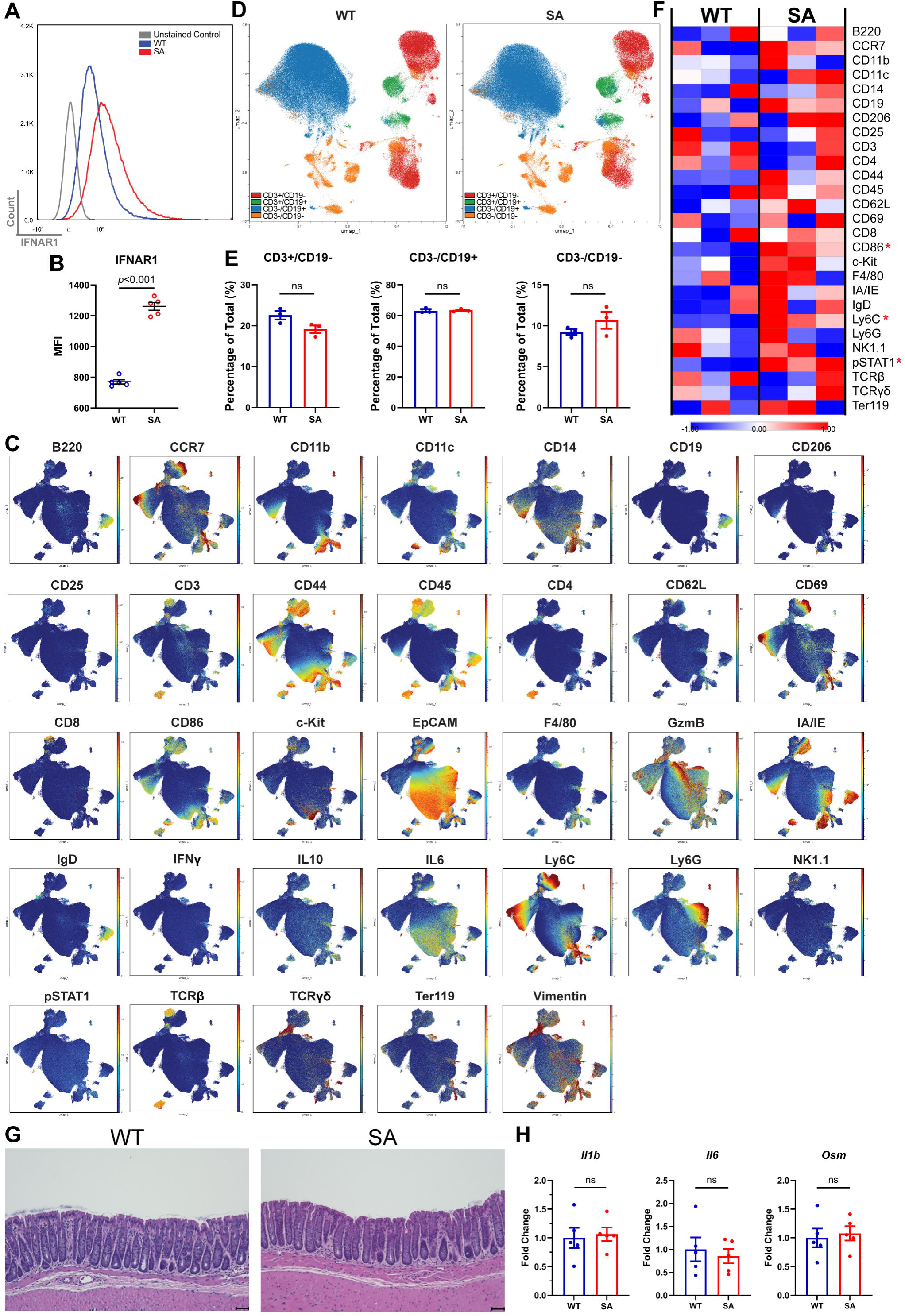
IFNAR1-SA mice exhibit higher surface expression of IFNAR1 and altered immune homeostasis in tissues. (A-B) Flow cytometry analysis and quantification of median fluorescence intensity (MFI) of IFNAR1 expression in spleen from WT (n=5) and SA (n=5) mice at baseline. **(C)** UMAP plots from mass cytometry showing expression of each target protein in the antibody panel (Supplemental Table 4) in mouse colon at baseline. **(D-E)** Mass cytometry analysis of spleen from WT (n=3) and SA (n=3) mice at baseline. **(D)** UMAP plot with cells colored by identity. **(E)** Proportions of immune cells classified by CD3 and CD19 positivity. ns: not significant. **(F)** Heatmap showing the mean expression of target proteins in the antibody panel (Supplemental Table 4), with those having *p* <0.05 by SAM (Significance Analysis of Microarrays) for their median expression indicated by red asterisks. **(G)** Representative H&E staining of colon from WT mice and SA mice at baseline. n=5 per group. Scale bar: 50 μm. **(H)** qRT-PCR validation of selected inflammatory cytokines in colonic biopsies. n=5 per group. ns: not significant.

**Supplemental Figure 3.**
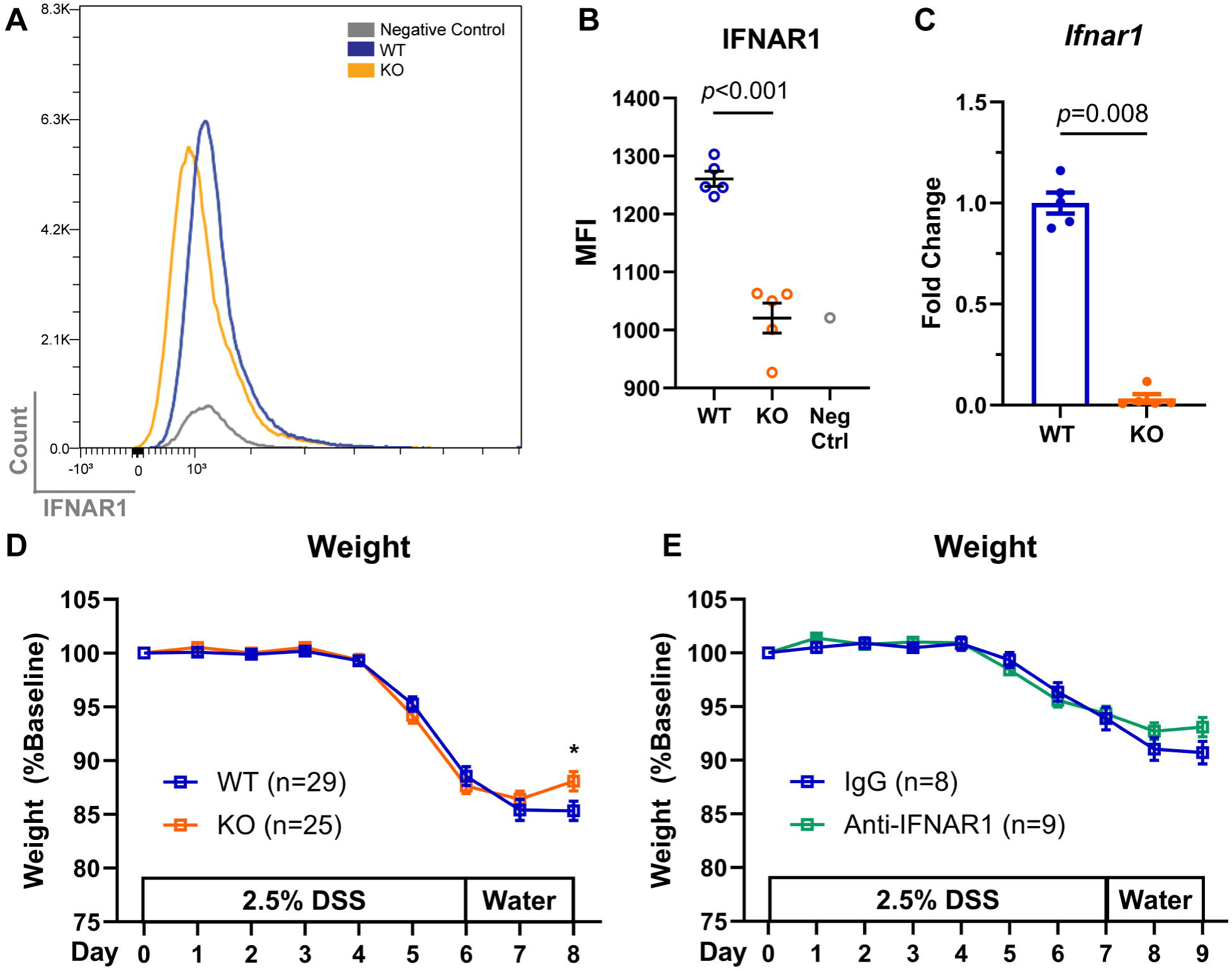
Post-natal genetic deletion or pharmacologic blocking of IFNAR1 attenuates DSS-induced colitis (A-C) Flow cytometry analysis **(A)** and quantification of median fluorescence intensity (MFI) **(B)** in spleen, and qRT-PCR analysis in colon **(C)** of IFNAR1 expression from WT (*Ifnar1*^fl/fl^, n=5) and KO (*Ifnar1*^fl/fl^; *Ubc*-CreERT2, n=5) mice following tamoxifen injection. **(D-E)** Weight change in DSS-induced colitis following tamoxifen-induced knockout of IFNAR1 **(D)** and administration of blocking anti-IFNAR1 antibody **(E)**.

**Supplemental Table 1.** Patient Charateristics.

| **Bulk RNA Sequencing** |  |  |  |
| --- | --- | --- | --- |
|  | **Non-IBD Control**<br>**(n=10)** | **Inactive UC**<br>**(n=6)** | **Active UC**<br>**(n=11)** |
| **Age (year)** | 56 ± 13 | 40 ± 16 | 41 ± 15 |
| **Male** | 6 (60%) | 4 (67%) | 7 (64%) |
| **Disease Extent** |  |  |  |
| Left-sided | N/A | 1 (17%) | 3 (27%) |
| Extensive | N/A | 5 (83%) | 8 (73%) |
| **Mayo Endoscopic Score** |  |  |  |
| Mayo 0 | N/A | 6 (100%) | 0 (0%) |
| Mayo 2 | N/A | 0 (0%) | 8 (73%) |
| Mayo 3 | N/A | 0 (0%) | 3 (27%) |
| **UC Therapy** |  |  |  |
| 5-ASA | N/A | 1 (17%) | 6 (55%) |
| Anti-TNF | N/A | 3 (50%) | 0 (0%) |
| Anti-Intergrin | N/A | 2 (33%) | 4 (36%) |
| None | N/A | 0 (0%) | 1 (9%) |
| **Immunofluorescence** |  |  |  |
|  | **Non-IBD Control**<br>**(n=5)** | **Active UC**<br>**(n=5)** |  |
| **Age (year)** | 50 ± 9 | 34 ± 15 |  |
| **Male** | 2 (40%) | 3 (60%) |  |
| **Disease Extent** |  |  |  |
| Left-sided | N/A | 1 (20%) |  |
| Extensive | N/A | 4 (80%) |  |
| **Mayo Endoscopic Score** |  |  |  |
| Mayo 2 | N/A | 1 (20%) |  |
| Mayo 3 | N/A | 4 (80%) |  |

Data are presented as mean ± SD or n (%). IBD: inflammatory bowel disease. UC: ulcerative colitis. 5-ASA: 5-aminosalicylic acid. TNF: tumor necrosis factor.

**Supplemental Table 2.** Interferome Analysis of IFN-I Signature Genes.

| Gene Name | Inducible by IFN Type |  |  |
| --- | --- | --- | --- |
| ACOD1 | I |  |  |
| AIM2 | I | II |  |
| BATF2 | I | II | III |
| BST2 | I | II | III |
| C1S | I | II |  |
| CASP1 | I | II |  |
| CD74 | I | II |  |
| CXCL10 | I | II |  |
| CXCL11 | I | II |  |
| FCAR | I | II |  |
| GATA3 |  | II |  |
| GBP2 | I | II |  |
| GBP4 | I | II |  |
| GMPR | I | II |  |
| HELZ2 | I | II | III |
| HLA-F | I | II |  |
| IFI16 | I | II |  |
| IFI30 | I | II |  |
| IFI44 | I | II |  |
| IFI44L | I | II |  |
| IFI6 | I | II | III |
| IFIT3 | I | II | III |
| IFITM1 | I | II |  |
| IFITM2 | I | II | III |
| IFITM3 | I | II | III |
| IFNAR2 | I | II |  |
| IRF1 | I | II | III |
| IRF4 | I | II |  |
| ISG15 | I | II | III |
| ISG20 | I | II |  |
| LAMP3 | I | II |  |
| LY6E | I | II |  |
| MNDA | I | II |  |
| MX2 | I | II |  |
| NCOA7 | I | II |  |
| OAS2 | I | II |  |
| OASL | I | II | III |
| PARP14 | I | II | III |
| PARP9 | I | II | III |
| PROCR | I | II |  |
| PSMB9 | I | II |  |
| PYHIN1 |  | II |  |
| RIPK2 | I | II |  |
| SAMD9L | I | II |  |
| SELL | I | II |  |
| SOCS1 | I | II |  |
| SOCS3 | I | II |  |
| STAT1 | I | II | III |
| STING1 | I | II |  |
| TAP1 | I | II | III |
| UBE2L6 | I | II | III |
| USP18 | I | II | III |
| WARS1 | I | II |  |
| XAF1 | I | II |  |

**Supplemental Table 3.** Antibodies for Immunofluoresence.

| **Target Protein** | **Manufacturer (Catalog Number)** | **Source** | **Dilution** |
| --- | --- | --- | --- |
| ISG15 | Santa Cruz (sc-166794) | Mouse Monoclonal | 1:150 |
| STAT1 | Cell Signaling Technology (9172) | Rabbit Polyclonal | 1:150 |
| Alexa Fluor 555 | Thermo Fisher Scientific (A21428) | Goat Anti-Rabbit | 1:500 |
| Alexa Fluor 488 | Thermo Fisher Scientific (A11029) | Goat Anti-Mouse | 1:500 |

**Supplemental Table 4.** Antibodies for Flow/Mass Cytometry.

| **Target Protein** | **Tag** | **Manufacturer (Catalog Number)** | **Application** |
| --- | --- | --- | --- |
| CD45 | Pacific Blue | BioLegend (157211) | Flow Cytometry |
| EpCAM | Alexa Fluor 700 | BioLegend (118239) | Flow Cytometry |
| IFNAR1 | APC | BioLegend (127313) | Flow Cytometry |
| B220 | 176Yb | Standard BioTools (3176002B) | Mass Cytometry |
| CCR7 | 164Er | Standard BioTools (3164013A) | Mass Cytometry |
| CD11b | 148Nd | Standard BioTools (3148003B) | Mass Cytometry |
| CD11c | 142Nd | Standard BioTools (3142003B) | Mass Cytometry |
| CD14 | 156Gd | Standard BioTools (3156009B) | Mass Cytometry |
| CD19 | 149Sm | Standard BioTools (3149002B) | Mass Cytometry |
| CD206 | 175Lu | BioLegend (151702) | Mass Cytometry |
| CD25 | 151Eu | Standard BioTools (3151007B) | Mass Cytometry |
| CD3 | 152Sm | Standard BioTools (3152004B) | Mass Cytometry |
| CD4 | 145Nd | Standard BioTools (3145002B) | Mass Cytometry |
| CD44 | 171Yb | Standard BioTools (3171003B) | Mass Cytometry |
| CD45 | 89Y | Standard BioTools (3089005B) | Mass Cytometry |
| CD62L | 160Gd | Standard BioTools (3160008B) | Mass Cytometry |
| CD69 | 143Nd | Standard BioTools (3143004B) | Mass Cytometry |
| CD69 | 161Dy | Standard BioTools (3143004B) | Mass Cytometry |
| CD8 | 116Cd | BioLegend (100755) | Mass Cytometry |
| CD86 | 172Yb | Standard BioTools (3172016B) | Mass Cytometry |
| c-Kit | 173Yb | Standard BioTools (3173004B) | Mass Cytometry |
| EpCAM | 166Er | Standard BioTools (3166014B) | Mass Cytometry |
| F4/80 | 146Nd | Standard BioTools (3146008B) | Mass Cytometry |
| GranzymeB | 198PT | Standard BioTools (3198002C) | Mass Cytometry |
| IA/IE | 209Bi | Standard BioTools (3209006B) | Mass Cytometry |
| IFNg | 165Ho | Standard BioTools (3165003B) | Mass Cytometry |
| IgD | 150Nd | Standard BioTools (3150011B) | Mass Cytometry |
| IL-10 | 158Gd | Standard BioTools (3158002C) | Mass Cytometry |
| IL-6 | 167Er | Standard BioTools (3167003C) | Mass Cytometry |
| Ly-6C | 162Dy | Standard BioTools (3162014B) | Mass Cytometry |
| Ly-6G | 141Pr | Standard BioTools (3141008B) | Mass Cytometry |
| NK1.1 | 170Er | Standard BioTools (3170002B) | Mass Cytometry |
| pSTAT1 | 153Eu | Standard BioTools (3153003A) | Mass Cytometry |
| TCRb | 169Tm | Standard BioTools (3169002B) | Mass Cytometry |
| TCRgd | 159Tb | Standard BioTools (3159012B) | Mass Cytometry |
| Ter119 | 154Sm | Standard BioTools (3154005B) | Mass Cytometry |
| Vimentin | 143ND | Standard BioTools (3143027D) | Mass Cytometry |

**Supplemental Table 5.**
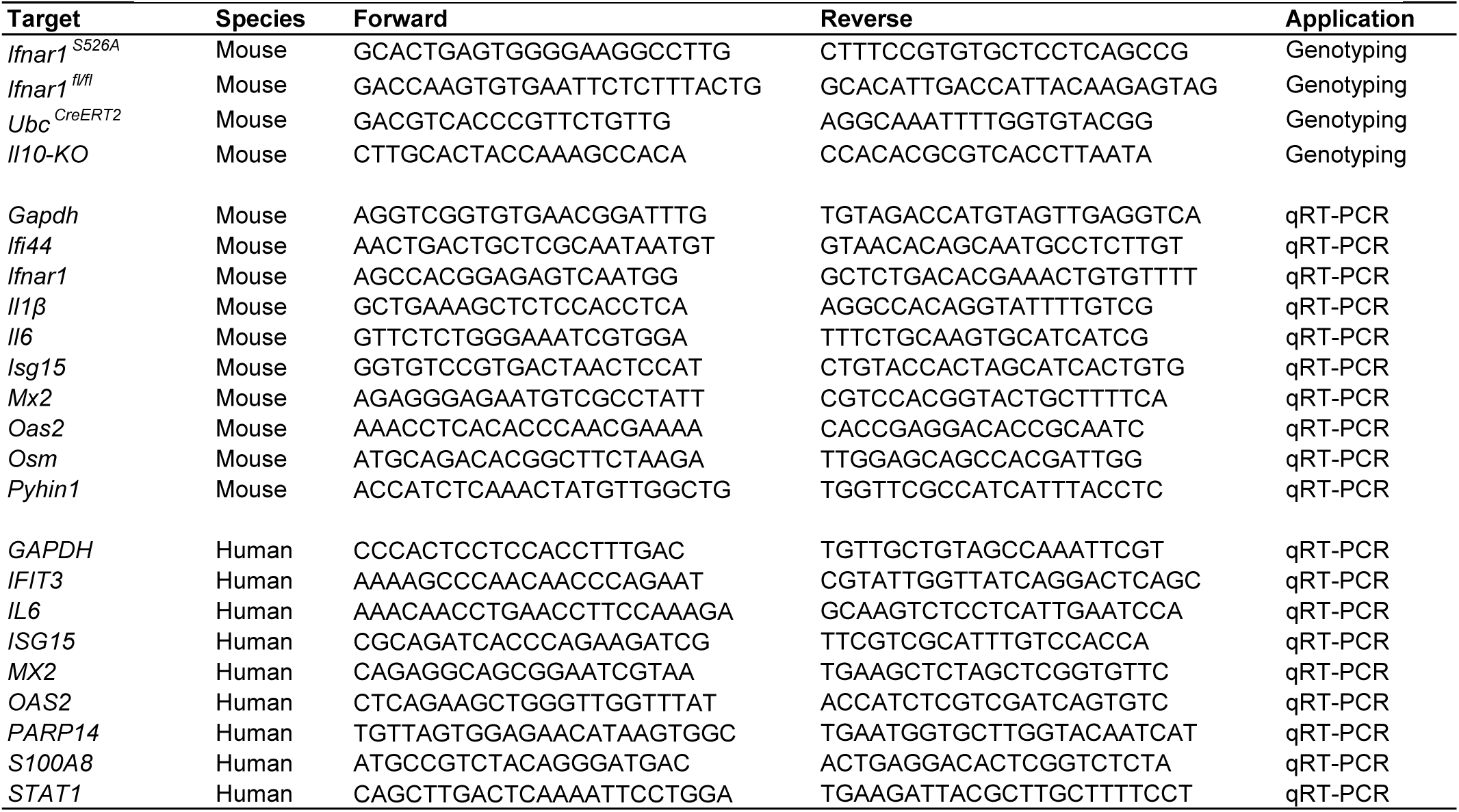
Primers for Genotyping and Quantitative Reverse Transcription PCR (qRT-PCR)

**Supplemental Table 6.** Assessment of Clinical Disease Activity Index (CDAI)

| **Score** | **Weight Loss** | **Stool Consistency** | **Blood** |
| --- | --- | --- | --- |
| 0 | None | Normal | None |
| 1 | 1-5% | Soft but still formed |  |
| 2 | 5-10% | Soft | Positive hemocult |
| 3 | 10-15% | Very soft and wet | Blood traces in stool |
| 4 | >15% | Watery | Gross rectal bleeding or bloody stool |

## REFERENCES

1. Boehmer D, and Zanoni I. Interferons in health and disease. Cell. 2025;188(17):4480–504.

2. Schneider WM, Chevillotte MD, and Rice CM. Interferon-stimulated genes: a complex web of host defenses. Annu Rev Immunol. 2014;32:513–45.

3. Ivashkiv LB, and Donlin LT. Regulation of type I interferon responses. Nat Rev Immunol. 2014;14(1):36–49.

4. Cohen RH, and Colgan SP. Mucosal Responses to Type II Interferon in IBD. Inflamm Bowel Dis. 2025.

5. Broggi A, Granucci F, and Zanoni I. Type III interferons: Balancing tissue tolerance and resistance to pathogen invasion. J Exp Med. 2020;217(1).

6. Psarras A, Wittmann M, and Vital EM. Emerging concepts of type I interferons in SLE pathogenesis and therapy. Nat Rev Rheumatol. 2022;18(10):575–90.

7. McNab F, Mayer-Barber K, Sher A, Wack A, and O’Garra A. Type I interferons in infectious disease. Nat Rev Immunol. 2015;15(2):87–103.

8. Chung H, Green PHR, Wang TC, and Kong XF. Interferon-Driven Immune Dysregulation in Down Syndrome: A Review of the Evidence. J Inflamm Res. 2021;14:5187–200.

9. Xavier RJ, and Podolsky DK. Unravelling the pathogenesis of inflammatory bowel disease. Nature. 2007;448(7152):427–34.

10. Friedrich M, Pohin M, and Powrie F. Cytokine Networks in the Pathophysiology of Inflammatory Bowel Disease. Immunity. 2019;50(4):992–1006.

11. Andreou NP, Legaki E, and Gazouli M. Inflammatory bowel disease pathobiology: the role of the interferon signature. Ann Gastroenterol. 2020;33(2):125–33.

12. Freedman MS, Coyle PK, Hellwig K, Singer B, Wynn D, Weinstock-Guttman B, et al. Twenty Years of Subcutaneous Interferon-Beta-1a for Multiple Sclerosis: Contemporary Perspectives. Neurol Ther. 2024;13(2):283–322.

13. Wang Y, MacDonald JK, Benchimol EI, Griffiths AM, Steinhart AH, Panaccione R, et al. Type I interferons for induction of remission in ulcerative colitis. Cochrane Database Syst Rev. 2015;2015(9):CD006790.

14. Mo Z, Tang J, Wu Z, Chen D, Xie D, and Wang P. Ulcerative colitis triggered by pegylated interferon alpha-2b in a patient with chronic hepatitis B: A case report and literature review. Liver Res. 2021;5(2):97–101.

15. Rodrigues S, Magro F, Soares J, Nunes AC, Lopes S, Marques M, et al. Case series: ulcerative colitis, multiple sclerosis, and interferon-beta 1a. Inflamm Bowel Dis. 2010;16(12):2001–3.

16. Tuna Y, Basar O, Dikici H, and Koklu S. Rapid onset of ulcerative colitis after treatment with interferon beta1a in a patient with multiple sclerosis. J Crohns Colitis. 2011;5(1):75–6.

17. Sifuentes-Dominguez L, Starokadomskyy P, Welch J, Gurram B, Park JY, Koduru P, et al. Mosaic Tetrasomy 9p Associated With Inflammatory Bowel Disease. J Crohns Colitis. 2019;13(11):1474–8.

18. Starokadomskyy P, Gemelli T, Rios JJ, Xing C, Wang RC, Li H, et al. DNA polymerase-alpha regulates the activation of type I interferons through cytosolic RNA:DNA synthesis. Nat Immunol. 2016;17(5):495–504.

19. McFarland AP, Savan R, Wagage S, Addison A, Ramakrishnan K, Karwan M, et al. Localized delivery of interferon-beta by Lactobacillus exacerbates experimental colitis. PloS one. 2011;6(2):e16967.

20. Katakura K, Lee J, Rachmilewitz D, Li G, Eckmann L, and Raz E. Toll-like receptor 9-induced type I IFN protects mice from experimental colitis. J Clin Invest. 2005;115(3):695–702.

21. Rauch I, Hainzl E, Rosebrock F, Heider S, Schwab C, Berry D, et al. Type I interferons have opposing effects during the emergence and recovery phases of colitis. Eur J Immunol. 2014;44(9):2749–60.

22. McElrath C, Espinosa V, Lin JD, Peng J, Sridhar R, Dutta O, et al. Critical role of interferons in gastrointestinal injury repair. Nat Commun. 2021;12(1):2624.

23. King MR, Kemp RA, and Gadeock S. Type I interferons in inflammatory bowel diseases: balancing barrier integrity, repair and inflammation in the intestinal epithelium. Front Med (Lausanne). 2025;12:1713424.

24. Salas A, Hernandez-Rocha C, Duijvestein M, Faubion W, McGovern D, Vermeire S, et al. JAK-STAT pathway targeting for the treatment of inflammatory bowel disease. Nat Rev Gastroenterol Hepatol. 2020;17(6):323–37.

25. Massimino L, Lamparelli LA, Houshyar Y, D’Alessio S, Peyrin-Biroulet L, Vetrano S, et al. The Inflammatory Bowel Disease Transcriptome and Metatranscriptome Meta-Analysis (IBD TaMMA) framework. Nat Comput Sci. 2021;1(8):511–5.

26. Rusinova I, Forster S, Yu S, Kannan A, Masse M, Cumming H, et al. Interferome v2.0: an updated database of annotated interferon-regulated genes. Nucleic Acids Res. 2013;41(Database issue):D1040–6.

27. Czarnewski P, Parigi SM, Sorini C, Diaz OE, Das S, Gagliani N, et al. Conserved transcriptomic profile between mouse and human colitis allows unsupervised patient stratification. Nat Commun. 2019;10(1):2892.

28. Holgersen K, Kvist PH, Markholst H, Hansen AK, and Holm TL. Characterisation of enterocolitis in the piroxicam-accelerated interleukin-10 knock out mouse — A model mimicking inflammatory bowel disease. J Crohns Colitis. 2014;8(2):147–60.

29. Garrido-Trigo A, Corraliza AM, Veny M, Dotti I, Melon-Ardanaz E, Rill A, et al. Macrophage and neutrophil heterogeneity at single-cell spatial resolution in human inflammatory bowel disease. Nat Commun. 2023;14(1):4506.

30. Uzzan M, Martin JC, Mesin L, Livanos AE, Castro-Dopico T, Huang R, et al. Ulcerative colitis is characterized by a plasmablast-skewed humoral response associated with disease activity. Nat Med. 2022;28(4):766–79.

31. Ho YT, Shimbo T, Wijaya E, Kitayama T, Takaki S, Ikegami K, et al. Longitudinal Single-Cell Transcriptomics Reveals a Role for Serpina3n-Mediated Resolution of Inflammation in a Mouse Colitis Model. Cell Mol Gastroenterol Hepatol. 2021;12(2):547–66.

32. Qian J, Zheng H, Huangfu WC, Liu J, Carbone CJ, Leu NA, et al. Pathogen recognition receptor signaling accelerates phosphorylation-dependent degradation of IFNAR1. PLoS Pathog. 2011;7(6):e1002065.

33. Bhattacharya S, Katlinski KV, Reichert M, Takano S, Brice A, Zhao B, et al. Triggering ubiquitination of IFNAR1 protects tissues from inflammatory injury. EMBO Mol Med. 2014;6(3):384–97.

34. Gui J, Gober M, Yang X, Katlinski KV, Marshall CM, Sharma M, et al. Therapeutic Elimination of the Type 1 Interferon Receptor for Treating Psoriatic Skin Inflammation. J Invest Dermatol. 2016;136(10):1990–2002.

35. Keubler LM, Buettner M, Hager C, and Bleich A. A Multihit Model: Colitis Lessons from the Interleukin-10-deficient Mouse. Inflamm Bowel Dis. 2015;21(8):1967–75.

36. Kotlarz D, Beier R, Murugan D, Diestelhorst J, Jensen O, Boztug K, et al. Loss of interleukin-10 signaling and infantile inflammatory bowel disease: implications for diagnosis and therapy. Gastroenterology. 2012;143(2):347–55.

37. Lee PY, Li Y, Kumagai Y, Xu Y, Weinstein JS, Kellner ES, et al. Type I interferon modulates monocyte recruitment and maturation in chronic inflammation. Am J Pathol. 2009;175(5):2023–33.

38. Huber JP, and Farrar JD. Regulation of effector and memory T-cell functions by type I interferon. Immunology. 2011;132(4):466–74.

39. Muskardin TLW, and Niewold TB. Type I interferon in rheumatic diseases. Nat Rev Rheumatol. 2018;14(4):214–28.

40. Rodrigues CP, Calçada RR, Arnaud M, Kaltenbach L, Manser M, Scheidegger A, et al. IFNAR1⁺ Neutrophils Orchestrate Chronic Inflammatory Damage Through Mitochondrial Remodeling. bioRxiv. 2025.

41. Aden K, Tran F, Ito G, Sheibani-Tezerji R, Lipinski S, Kuiper JW, et al. ATG16L1 orchestrates interleukin-22 signaling in the intestinal epithelium via cGAS-STING. J Exp Med. 2018;215(11):2868–86.

42. Salazar-Mather TP, Lewis CA, and Biron CA. Type I interferons regulate inflammatory cell trafficking and macrophage inflammatory protein 1alpha delivery to the liver. J Clin Invest. 2002;110(3):321–30.

43. Majer O, Bourgeois C, Zwolanek F, Lassnig C, Kerjaschki D, Mack M, et al. Type I interferons promote fatal immunopathology by regulating inflammatory monocytes and neutrophils during Candida infections. PLoS Pathog. 2012;8(7):e1002811.

44. Diamond MS, Kinder M, Matsushita H, Mashayekhi M, Dunn GP, Archambault JM, et al. Type I interferon is selectively required by dendritic cells for immune rejection of tumors. J Exp Med. 2011;208(10):1989–2003.

45. Jarry A, Malard F, Bou-Hanna C, Meurette G, Mohty M, Mosnier JF, et al. Interferon-Alpha Promotes Th1 Response and Epithelial Apoptosis via Inflammasome Activation in Human Intestinal Mucosa. Cell Mol Gastroenterol Hepatol. 2017;3(1):72–81.

46. Jostins L, Ripke S, Weersma RK, Duerr RH, McGovern DP, Hui KY, et al. Host-microbe interactions have shaped the genetic architecture of inflammatory bowel disease. Nature. 2012;491(7422):119–24.

47. Horesh ME, Martin-Fernandez M, Gruber C, Buta S, Le Voyer T, Puzenat E, et al. Individuals with JAK1 variants are affected by syndromic features encompassing autoimmunity, atopy, colitis, and dermatitis. J Exp Med. 2024;221(6).

48. Legeret C, Meyer BJ, Rovina A, Deigendesch N, Berger CT, Daikeler T, et al. JAK Inhibition in a Patient with X-Linked Reticulate Pigmentary Disorder. J Clin Immunol. 2021;41(1):212–6.

49. Samie M, Lim J, Verschueren E, Baughman JM, Peng I, Wong A, et al. Selective autophagy of the adaptor TRIF regulates innate inflammatory signaling. Nat Immunol. 2018;19(3):246–54.

50. Mavragani CP, Nezos A, Dovrolis N, Andreou NP, Legaki E, Sechi LA, et al. Type I and II Interferon Signatures Can Predict the Response to Anti-TNF Agents in Inflammatory Bowel Disease Patients: Involvement of the Microbiota. Inflamm Bowel Dis. 2020;26(10):1543–53.

51. Ostvik AE, Svendsen TD, Granlund AVB, Doseth B, Skovdahl HK, Bakke I, et al. Intestinal Epithelial Cells Express Immunomodulatory ISG15 During Active Ulcerative Colitis and Crohn’s Disease. J Crohns Colitis. 2020;14(7):920–34.

52. Flood P, Fanning A, Woznicki JA, Crowley T, Christopher A, Vaccaro A, et al. DNA sensor-associated type I interferon signaling is increased in ulcerative colitis and induces JAK-dependent inflammatory cell death in colonic organoids. Am J Physiol Gastrointest Liver Physiol. 2022;323(5):G439–G60.

53. Yin J, El-Najjar Y, Cordova N, Touma MJ, Nguyen N, Boktor M, et al. Short-Term Use of Upadacitinib in Combination With Biologic Therapy for Inducing Clinical Remission in Patients With Active Inflammatory Bowel Disease. Inflamm Bowel Dis. 2024;30(10):1914–6.

54. Honap S, Agorogianni A, Colwill MJ, Mehta SK, Donovan F, Pollok R, et al. JAK inhibitors for inflammatory bowel disease: recent advances. Frontline Gastroenterol. 2024;15(1):59–69.

55. Danese S, and Peyrin-Biroulet L. Selective Tyrosine Kinase 2 Inhibition for Treatment of Inflammatory Bowel Disease: New Hope on the Rise. Inflamm Bowel Dis. 2021;27(12):2023–30.

56. Zeeff SB, Kunne C, Bouma G, de Vries RB, and Te Velde AA. Actual Usage and Quality of Experimental Colitis Models in Preclinical Efficacy Testing: A Scoping Review. Inflamm Bowel Dis. 2016;22(6):1296–305.

57. Hale LP, Gottfried MR, and Swidsinski A. Piroxicam treatment of IL-10-deficient mice enhances colonic epithelial apoptosis and mucosal exposure to intestinal bacteria. Inflamm Bowel Dis. 2005;11(12):1060–9.

58. Wirtz S, Popp V, Kindermann M, Gerlach K, Weigmann B, Fichtner-Feigl S, et al. Chemically induced mouse models of acute and chronic intestinal inflammation. Nat Protoc. 2017;12(7):1295–309.

